# Urban-use pesticides in stormwater ponds and their accumulation in biofilms

**DOI:** 10.1101/2023.11.28.568083

**Authors:** Gab Izma, Melanie Raby, Ryan Prosser, Rebecca Rooney

**Author notes:** corresponding author (Dr. Rebecca Rooney, 200 University Ave W. Waterloo, Ontario, Canada, N2L 3G1).

## Abstract

Stormwater ponds frequently receive urban runoff, increasing the likelihood of pesticide contamination. Biofilms growing in surface waters of these ponds are known to accumulate a range of aquatic contaminants, paradoxically providing both water purification services and potentially posing a threat to urban wildlife. Thus, sampling biofilms in stormwater ponds may be a critical and biologically relevant tool for characterizing pesticide contamination and toxicity in urban environments. Here, we aimed to investigate pesticide occurrences at 21 stormwater ponds in Brampton, ON, one of Canada’s fastest growing municipalities, and quantify their accumulation in biofilm. Over nine weeks, we collected time-integrated composite water and biofilm samples for analysis of ∼500 current-use and legacy pesticides. Thirty-two pesticide compounds were detected across both matrices, with 2,4-D, MCPA, MCPP, azoxystrobin, bentazon, triclopyr, and diuron having near-ubiquitous occurrences. Several compounds not typically monitored in pesticide suites (e.g., melamine and nicotine) were also detected, but only in biofilms. Overall, 56% of analytes detected in biofilms were not found in water samples, indicating traditional pesticide monitoring practices fail to capture all exposure routes, as even when pesticides are below detection levels in water, organisms may still be exposed via dietary pathways. Calculated bioconcentration factors ranged from 4.2 – 1275 and were not predicted by standard pesticide physicochemical properties. Monitoring biofilms provides a sensitive and comprehensive supplement to water sampling for pesticide quantification in urban areas, and identifying pesticide occurrences in stormwater could improve source-tracking efforts in the future. Further research is needed to understand the mechanisms driving pesticide accumulation, to investigate toxicity risks associated with pesticide-contaminated biofilm, and to evaluate whether pesticide accumulation in stormwater pond biofilms represents a route through which contaminants are mobilized into the surrounding terrestrial and downstream aquatic environments.

## 1. Introduction

In urban areas, stormwater management ponds are employed to accommodate hydrological alterations imposed by urbanization and sprawl. Runoff accumulates in these retention ponds via stormwater drainage systems and can be loaded with sediment and contaminants (Göbel et al., 2007), including road salt, heavy metals (Pitt et al., 1995; Zgheib et al., 2012), pesticides (Burant et al., 2018), oil (Brown et al., 1985), and fecal bacteria (Valenca et al., 2022). Discharge from stormwater ponds is an important source of chemical and thermal pollution into freshwater systems, with downstream consequences, such as habitat degradation and biodiversity loss, being well documented (e.g., Hatt et al., 2004; Burns et al., 2012; Park & Roesner, 2012; Zgheib et al., 2012). Frequently overlooked, however, are the ecological implications of these stressors within the ponds, which have become urban habitat for aquatic wildlife (Le Viol et al., 2009; Perron et al., 2020).

Although stormwater ponds primarily serve flood mitigation objectives, they are also frequently designed to improve water quality, in recognition of their role in transforming and accumulating contaminants to protect downstream waters (e.g., MOE, 2003; EPA, 2009). However, stormwater ponds are not routinely monitored for contaminants. In particular, levels of current-use and legacy pesticides in stormwater ponds are not well documented; for example, the only other evaluation of pesticide levels in stormwater ponds in Ontario was published in 1994 by Struger et al. A stormwater pond study was conducted in 2000 by Bishop et al., but they only looked at pesticides present in sediment, not water. This knowledge gap is emphasized in a critical review by Chen et al. (2019) that covered pesticides in stormwater and identified an urgent need for research quantifying a wider scope of pesticide mixtures and investigating the biological and physicochemical drivers of pesticide occurrence variability in urban runoff. Urban-use pesticides, such as methylchlorophenoxypropionic acid (MCPP) and 2,4-dichlorophenoxyacetic acid (2,4-D), accumulate in stormwater ponds (Struger et al., 1994; Raina et al., 2011; Chen et al., 2019; Flanagan et al., 2021) and could present toxicological risks to urban wildlife residing in stormwater ponds (Allinson et al., 2015; Flanagan et al., 2021). Moreover, contaminant retention within the ponds can be compromised, particularly during flood events when pond sediments are disturbed (Vulava et al., 2019; Spahr et al., 2020). Sediment resuspension can cause remobilization of contaminants and threaten downstream ecosystems. The lack of routine monitoring in stormwater ponds complicates efforts to predict both within-pond risks to biota, and the potential for pesticide remobilization into surrounding aquatic ecosystems.

Pesticides can enter stormwater in urban environments through a variety of uses, including construction activities, personal and pet care (e.g., bed bug or flea treatments), household cleaning, and municipal landscaping (Meftaul et al., 2020; Spahr et al., 2020; OMAFRA, 2021). Pesticides may also be applied directly to stormwater ponds, for example, methoprene is a larvicide used to kill mosquitos and mitigate the public health risk of West Nile Virus in some stormwater ponds in Ontario (Peel Public Health, personal communication, October 28, 2022). The persistence and accumulation of pesticides in aquatic environments is highly dependent on their physiochemical properties, such as their solubility in water, which influences the partitioning of pesticides into water, sediment, or organic material (Gramatica & Di Guardo, 2002). Thus, traditional assessment using water sampling techniques may fail to capture pesticides in biologically relevant material, such as biofilms (Rooney et al., 2020; Mahler et al., 2020; Bonnineau et al., 2021). Recent evidence suggests that biofilms can accumulate a range of contaminants, including pesticides (Mahler et al., 2020; Fernandez et al., 2020; Beecraft & Rooney, 2021; Ijzerman et al., 2023). Beecraft and Rooney (2021) found glyphosate to be two to four times higher in biofilms compared with surrounding water, while Lawrence et al. (2001) proposed that biofilms are sinks for herbicides after discovering that diclofop methyl and atrazine were detected in river biofilm samples despite not being detected in water samples. Most recently, Rheinheimer Dos Santos et al. (2023) found pesticide occurrences were underrepresented in grab water samples compared to epilithic biofilm collected from natural substrates in both agricultural and urban stream systems. Primary production and decomposition in aquatic ecosystems are driven by microbial activity (Hall & Meyer, 1998; Zou et al., 2016), thus biofilms are key energy sources for many invertebrate and fish species but may represent trophic links through which contaminants can be passed (Bonnineau et al., 2021). Not only do biofilms loaded with pesticides potentially pose a toxic threat to biota through dietary transfer, but connections between aquatic and terrestrial environments (e.g., during emergence of adult stage invertebrates or amphibians) could facilitate mobilization of pesticides from stormwater systems into neighboring terrestrial habitats. Thus, it is critical to understand contamination in both stormwater pond water and biofilm matrices and to measure the magnitude of accumulation within biofilms.

With a high surface area to volume ratio, biofilms can be exposed to contaminants from both the water and the surfaces to which they attach. Pesticide accumulation is thought to occur via sorption and passive diffusion (Flemming 1995; Chaumet et al., 2019b), where the hydrophobicity of a pesticide has been previously hypothesized as the key determinant of uptake and accumulation within biofilms. For example, Nikkila et al. (2001) monitored atrazine uptake and depuration by a river biofilm and found that atrazine was rapidly taken up by periphyton but was minimally bioconcentrated due to low hydrophobicity. Capacity for pesticide accumulation in biofilms may also be influenced by surrounding water quality parameters, such as salinization, which is known to lower diatom diversity and composition (Ziemann et al., 2001; Coring & Bathe, 2011; Van Meter et al., 2011), microbial respiration in heterotrophic biofilms (Entrekin et al., 2019; Martinez et al., 2020), and photosynthetic rates in autotrophic biofilms (Cook & Francoeur, 2013). Although environmental drivers of pesticide accumulation remain understudied, changes resulting in altered biofilm community structures, and thus their lipid content, may influence biofilms’ capacity to accumulate organic chemicals (Wang et al., 1999).

In this study, our objectives were to: 1) determine the maximum concentration and occurrence of herbicides, insecticides, and fungicides in stormwater pond water and in biofilms, and compare those concentrations to exposure thresholds to assess potential risk to aquatic biota; 2) evaluate the relationship between pesticide occurrences in each matrix and the physicochemical properties of the pesticides; 3) compare the concentrations of pesticides in water to those in cultivated biofilm to quantify accumulation using bioconcentration factors; and 4) investigate the relationship between bioconcentration factors and the physicochemical properties of the pesticides we detected alongside the general quality of the water in the stormwater ponds and the chlorophyll-*a* content and ash-free dry weight of the biofilms. We hypothesized that the accumulation of pesticides into biofilm would be related to the physicochemical properties of the pesticides, and that pesticides that are more soluble in water will accumulate in biofilm the least and be detected more frequently in the water matrix. We also hypothesized that pesticides with higher octanol-water (Log K_ow_) or soil adsorption (Log K_oc_) coefficients will accumulate in biofilm more and be more frequently detected in biofilm than in water.

## 2. Material and methods

### 2.1 Location of study

The City of Brampton, Ontario, is one of Canada’s fastest growing municipalities and experiences high rates of development and urban expansion (Statistics Canada 2022; City of Brampton 2022). The resulting impervious cover totals 7,888 ha, or 45.4%, of the City’s area (TRCA, n.d.). Located within two major watersheds, the West Humber River watershed and the Credit River watershed, Brampton is home to more than 180 stormwater ponds that ultimately feed into Lake Ontario (City of Brampton, n.d.). Twenty-one of these stormwater ponds were selected for this study (Figure 1). The impervious cover within a 300 m buffer surrounding each stormwater pond ranged from 10.9 to 55.2% (range = 69,630 – 247,521 m^2^, mean = 159,060 m^2^, standard deviation = 45,224 m^2^) to yield a gradient in urbanization intensity (McIsaac 2022). The 21 stormwater ponds drain into either Etobicoke Creek, Mimico Creek, the Credit River, or the Humber River. As wet ponds, these locations are designed and managed to retain water all year long. The ponds ranged in size from 1051 to 3686 m^2^ (mean = 2272 m^2^, standard deviation = 851 m^2^), were a minimum of 10 years old, and had no history of dredging within the past 10 years.

**Figure 1.**
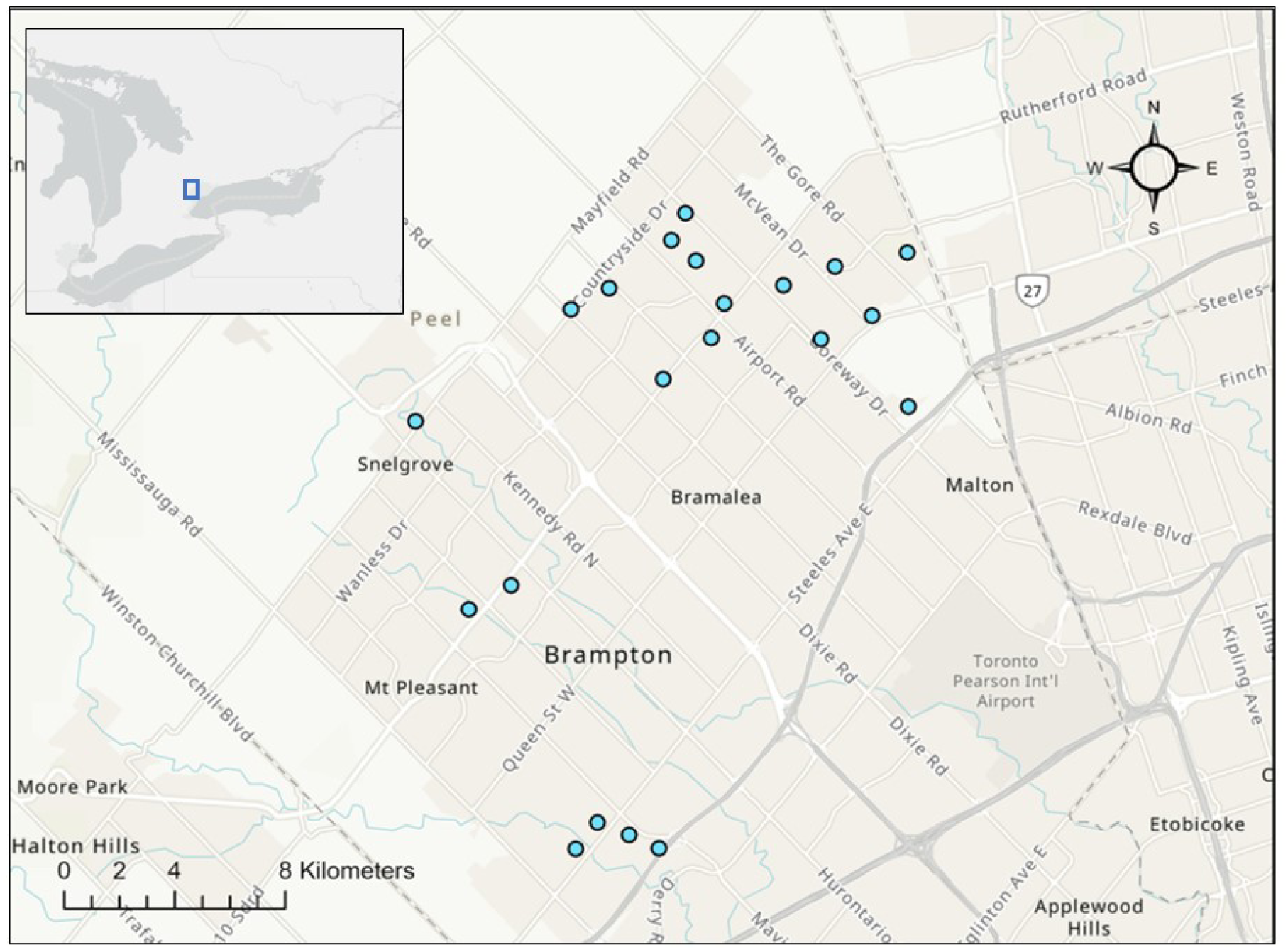
Map of stormwater pond sites (blue circles) in Brampton, Ontario. Map created in ARCGIS on July 5th, 2023.

### 2.2 Collection and analysis of water samples

We collected a time-integrated stormwater surface sample to characterize pesticide concentrations in each pond over a nine-week study period between May 25^th^ and July 25^th^, 2022. This was achieved by adding a 150 mL weekly subsample, taken at ∼20 cm depth, to a 1.35 L composite that we kept at –20 °C until submission to the Agriculture and Food Laboratory (AFL) (ISO/IEC 17025 accredited) in Guelph, Ontario, for a comprehensive analysis of ∼500 current-use and legacy pesticides using multi-residue liquid chromatography/electrospray ionization-tandem mass spectrometry (LC/ESI-MS/MS) and gas chromatography-tandem mass spectrometry (GC-MS/MS). Details of this analysis, including lists of analytes and their detection and quantification limits are provided in the Supplemental Materials section.

We measured water quality regularly over the same nine-week period (Table 1). At each stormwater pond, at 3 points in the open water and ∼ 20 cm deep, we measured water temperature (°C) and conductivity (mS cm^-1^), using a Hach HQ1140 Portable Conductivity/TDS Meter (Hach Sales and Services L.P., London, Canada) and we measured dissolved oxygen (DO; mg L^-1^), using a YSI ProSolo ODO Optical Dissolved Oxygen Meter (Xylem Inc., Miami, FL).

**Table 1.** Water chemistry parameters measured at each stormwater pond site (n = 21) in Brampton, Ontario. WWL = Waterloo Wetland Lab; ALS = ALS Laboratories; AFL = University of Guelph Agriculture and Food Laboratory.

| Parameter | Location of analysis | Method | Frequency of analysis |
| --- | --- | --- | --- |
| Surface water temperature | In-situ | Hach multi-probe meter | Weekly |
| Conductivity | In-situ | Hach multi-probe meter | Weekly |
| DO | In-situ | Hach multi-probe meter | Weekly |
| pH | WWL | Thermo-Orion pH probe and benchtop meter | Weekly |
| Chloride | WWL | Colorimetric | Weekly |
| Chlorophyll-*a* | WWL | Fluorometric | June and July |
| TSS | WWL | Gravimetric | Weekly |
| Ammonia | ALS | Fluorometric | Once |
| Nitrate | ALS | Ion chromatography | Once |
| Nitrite | ALS | Ion chromatography | Once |
| Orthophosphate | ALS | Colorimetric | Once |
| *E. coli* and coliforms | AFL | Plate counts | Once |
| Pesticides | AFL | Multi-residue LC-MS | Once* |

\*analysis completed on time-integrated composite of weekly water samples.

Each week, we combined 50 mL surface water grab samples from ∼20 cm depth within a high-density polyethylene Nalgene bottle to make an additional time-integrated composite for the purpose of ex situ measurements. We kept the weekly water sample on ice during transport to the Waterloo Wetland Lab (WWL) in Waterloo, Ontario, where we determined total suspended solids (TSS; mg L^-1^) gravimetrically, and measured pH using an Orion 9156BNWP Combination pH Electrode and Orion Star A211 Benchtop pH Meter (Thermo Fisher Scientific, Waltham, MA). We measured chloride ion concentration (Cl^-^; mg L^-1^) colorimetrically, using a Chloride Checker® HC (Hannah Instruments, Woonsocket, RI). Once in June (week of June 21^st^, 2022) and once in July (week of July 11^th^, 2022), we collected additional composite water grab samples for the fluorometric determination of chlorophyll-*a (*ug L^-1^) using a Turner Designs TD 10AU fluorometer (Turner Designs Inc., Sunnyvale, California) at the WWL. This machine is calibrated annually using a calibration curve which was created from a dilution series of chlorophyll-*a* standard and validated with a Biochrom Ultrospec 3100 spectrophotometer (Biochrom Ltd, Cambridge, UK).

From June 21^st^ – 24^th^, 2022, we collected a water grab sample from each site and submitted them for nutrient analysis at ALS laboratories in Waterloo, Ontario, which included fluorometric determination of ammonia (NH_3_), ion chromatographic determination of nitrate (NO_3_) and nitrite (NO_2_), and calorimetric analysis of orthophosphate. Finally, following a precipitation event on June 7^th^, 2022, we collected and submitted surface grab water samples from each stormwater pond to the Agriculture and Food Laboratory (AFL) in Guelph, Ontario, for plate-count analysis of *E. coli* and total coliform bacteria.

We used the Canadian Council of Ministers of the Environment (CCME) Water Quality Index (WQI) Calculator version 1.2 to calculate WQI scores for each stormwater pond site. The following 13 parameters were included in this calculation: pH, conductivity (mS cm^-1^), DO (mg L^-1^), surface water temperature (°C), chloride concentration (mg L^-1^), chlorophyll-*a* concentration (µg L^-1^), TSS (mg L^-1^), ammonia (mg L^-1^), nitrate (mg L^-1^), nitrite (mg L^-1^), phosphorus (mg L^-1^), coliforms (cfu per 100 mL), and *E. coli* (cfu per 100 mL). The Canadian Environmental Quality Guidelines for the protection of aquatic life reported by CCME (accessed at: https://ccme.ca/en/resources#) were prioritized for these calculations, and where none existed, we used guidelines from the Ontario Provincial Water Quality Objectives (accessed at: https://www.ontario.ca/page/water-management-policies-guidelines-provincial-water-quality-objectives#section-3), the USEPA Water Quality Criteria for Aquatic Life (accessed at: https://www.epa.gov/wqc/national-recommended-water-quality-criteria-aquatic-life-criteria-table), and the USEPA Aquatic Life Benchmarks (https://www.epa.gov/pesticide-science-and-assessing-pesticide-risks/aquatic-life-benchmarks-and-ecological-risk#aquatic-benchmarks). Table S1 shows the WQI scores for each stormwater pond site.

### 2.3 Collection and analysis of biofilm samples

To culture biofilm *in-situ*, we deployed ten standard substrates (acrylic plates 20.2 by 44.4 cm in area) in each of the 21 stormwater ponds, following the deployment protocol described by Rooney et al. (2020). In brief, we acid-bathed the plates, rinsed them three times with deionized water, and gently scuffed them with fine sandpaper prior to deploying them on floating racks (Figure S1. A) that suspended them ∼10 cm beneath the water’s surface (Figure S1. B). We anchored the racks at depths between 60 and 100 cm in areas clear of vegetation and shading. After 54 days of deployment, we retrieved the plates and harvested the biofilm by scraping each side of each plate into a composite sample for each sampler rack, and quantitatively transferring the resulting biofilm slurry using a measured amount of distilled water to rinse.

We removed an aliquot of 300 – 500 g (wet weight) from each composite biofilm sample and freeze-dried it, using a FreeZone I Labconco benchtop freeze dryer (Labconco Corporation, Kansas City, Missouri). We then submitted the freeze-dried biofilm to AFL for analysis of the same suite of pesticides as described for water samples. Of note, two sites did not produce adequate biofilm mass for all pesticide analysis screens and were only analyzed for phenoxy herbicides and polar pesticides (please see SI for a list of analytes). However, the other sites (n = 19) had sufficient biofilm mass to complete all pesticide screens.

For each biofilm sample, we measured dry mass and ash-free dry weight (AFDW) gravimetrically using a Thermo Scientific F6010 Thermolyne furnace (Thermo Fisher Scientific Inc., Asheville, North Carolina). To measure chlorophyll-*a* levels in the biofilm fluorometrically, we sonicated the samples at 90% amplitude for three 30-second intervals and centrifuged at 3500 rpm for 6 minutes prior to fluorometric assessment using the same method applied to the water samples.

### 2.5 Characterization of pesticides

We used databases such as the Pesticide Properties Database (PPDB; accessed at: https://sitem.herts.ac.uk/aeru/ppdb/en/index.htm), and the Pesticide Action Network Pesticide Database (http://www.pesticideinfo.org/) to extract information regarding the chemical structures and properties of the pesticides detected in our samples. We supplemented these sources with chemical profiles reported by PubChem online database (accessed at: https://pubchem.ncbi.nlm.nih.gov/) and ChemSpider online database (accessed at: https://www.chemspider.com/Default.aspx). Specifically, we collected each chemical’s CAS number, chemical class, mode of action, octanol-water partition coefficient (Log K_ow_), solubility in water, soil adsorption coefficient (Log K_oc_), persistence in aquatic environments, and potential sources. We also used Health Canada’s Pesticide Product Information Database (PPID; accessed at: https://pest-control.canada.ca/pesticide-registry/en/product-search.html) to determine the status of pesticide permits.

### 2.6 Data analysis

#### Pesticide occurrences and concentrations and comparison to toxicity thresholds

We visualized the relative occurrence of pesticide types and their detections in water vs biofilm (objective 1) using Microsoft Excel. To support comparison to previous studies (e.g., Struger et al., 1994; Rooney et al. 2020), we grouped detected pesticides into four categories based on type: herbicides, insecticides, fungicides, and other biocides. Some pesticides bridged more than one type (e.g., nicotine is historically associated with insecticidal uses, but present-day detections are more likely with the result of improper cigarette disposal), and in these cases we selected the most probable type given the urban setting.

To facilitate comparisons between water and biofilm samples, we report pesticide concentrations as parts per billion (ppb), rather than as µg L^-1^ in water samples and µg kg^-1^ in biofilm samples. To estimate potential risks posed to stormwater organisms, the levels of pesticides detected in the stormwater were compared to known levels of concern for aquatic exposures provided by the Canadian Council of Ministers of the Environment (CCME) and the Pesticide Properties Database (PPDB).

#### Pesticide detections and their physicochemical properties

We tested whether the number of detections of pesticides in water and biofilm matrices depended on the pesticide properties (Log K_ow_, Log K_oc_, water solubility; objective 2) using generalized linear models with a Poisson distribution and log link function, because of its suitability for count data. These properties were selected because of their known influence on chemical fate in the environment (Gramatica & Di Guardo, 2002). As active uptake of pesticides by biofilms is unlikely, we hypothesized accumulation would be dependent on each chemical’s inclination for sorption and absorption on organic surfaces, in addition to its concentration in the water.

#### Bioconcentration factor calculations

The accumulation of heavy metals, pharmaceuticals, and pesticides from the surrounding water and sediment into biofilm has been characterized as bioconcentration by other researchers (Ancion et al., 2014; Bonnineau et al., 2021; Rooney et al., 2020). This characterization is pragmatic: microbial organisms in biofilms are not analyzed individually to measure bioconcentration by each microbe, but biofilm is consumed whole by many aquatic organisms. Thus, the increase in contaminant concentration from water and sediment to biofilm creates an exposure magnification that is analogous to bioconcentration from an ecotoxicology perspective. However, because it is not certain whether pesticides adsorb onto the extracellular polymeric substance (EPS) matrix or to the surface of microbial cells, or whether pesticides enter microbial organisms through active or passive transport across cell membranes, the general term “accumulation” is more appropriately used over “bioconcentration” or “bioaccumulation.”. In this study we use the calculation of bioconcentration factors (BCFs) as useful indices of pesticide accumulation, but we do not imply that pesticides enter the cells of microorganisms in the biofilms.

Bioconcentration factors (BCFs) for each pesticide at each site were calculated using the pesticide concentrations reported in both the stormwater and in the freeze-dried biofilm by AFL (objective 3). We used the calculation:

BCF = (*ppb* pesticides in freeze-dried biofilm)/(*ppb* pesticides in stormwater)

as described by Beecraft and Rooney (2021) and Rooney et al. (2020). We divided our reported BCFs into two groups: calculated and estimated BCFs, where calculated BCFs represent a more confident assessment in which biofilm pesticide concentrations were measured above the method quantification limit (MQL) in both the biofilm and the water from the stormwater ponds. If a pesticide was quantified in a biofilm sample, but not in the surrounding stormwater, we used the method detection limit (MDL) value for the water to estimate a BCF that was as conservative as possible. Estimated BCFs represent instances where concentrations in the biofilm were below the MQL but above the MDL, in which case the MDL limit value was used. Both estimated and calculated BCFs were reported, however only calculated BCFs were averaged and used in statistical analyses.

#### Bioconcentrations factors and pesticide physicochemical properties

We used generalized linear models to investigate the relationship between calculated bioconcentration factors (BCFs) and Log K_ow_, Log K_oc_, and water solubility of the pesticides, water quality in each stormwater pond, and chlorophyll-*a* content and AFDW of biofilm (objective 4). These generalized linear models used a gamma distribution with a log link function, chosen for its suitability in accommodating the positively skewed BCF data. An average calculated BCF was used for pesticides with more than one calculated BCF.

We used R Statistical Software (version 4.2.2.) in RStudio (version 2023.06.1; Rstudio Team 2021) for all analyses. To implement generalized linear models, we used the glm function in base R. We evaluated model fit for all the generalized linear models by comparing the null deviance to the residual deviance and assessing statistical significance of each model term.

## 3. Results and Discussion

### 3.1 Concentration and occurrence of pesticides in stormwater and biofilm

We found 32 pesticides across our 21 sampling sites and matrices. The minimum, maximum, and mean concentrations of each pesticide observed are reported in Table 2. Pesticide contamination is clearly widespread in urban areas. For example, the herbicides 2,4-D, MCPA, MCPP, triclopyr, and bentazon were nearly ubiquitous in water samples. Neonicotinoids were also frequently found in water samples, with detectable levels of clothianidin and imidacloprid at 67% and 62% of sites, respectively. Despite provincial legislation introduced in 2015 to reduce their use by 80% (OMECP, 2015), neonicotinoids still have both urban and agricultural applications (e.g., in lawn care and domestic pest management; PMRA 2016; OMAFRA 2017; Batikian et al., 2019) and are evidently as common in urban stormwater as in Ontario’s agricultural streams (Raby et al., 2022). Fungicides were less commonly detected in stormwater, but by sampling biofilms we were able to determine that they are also widespread in urban areas. For example, biofilm samples had detectable levels of azoxystrobin and thiabendazole at 95% and 81% of sampling sites, respectively. Biofilm sampling also revealed the widespread presence of nicotine in 52% of the sites sampled.

**Table 2.**
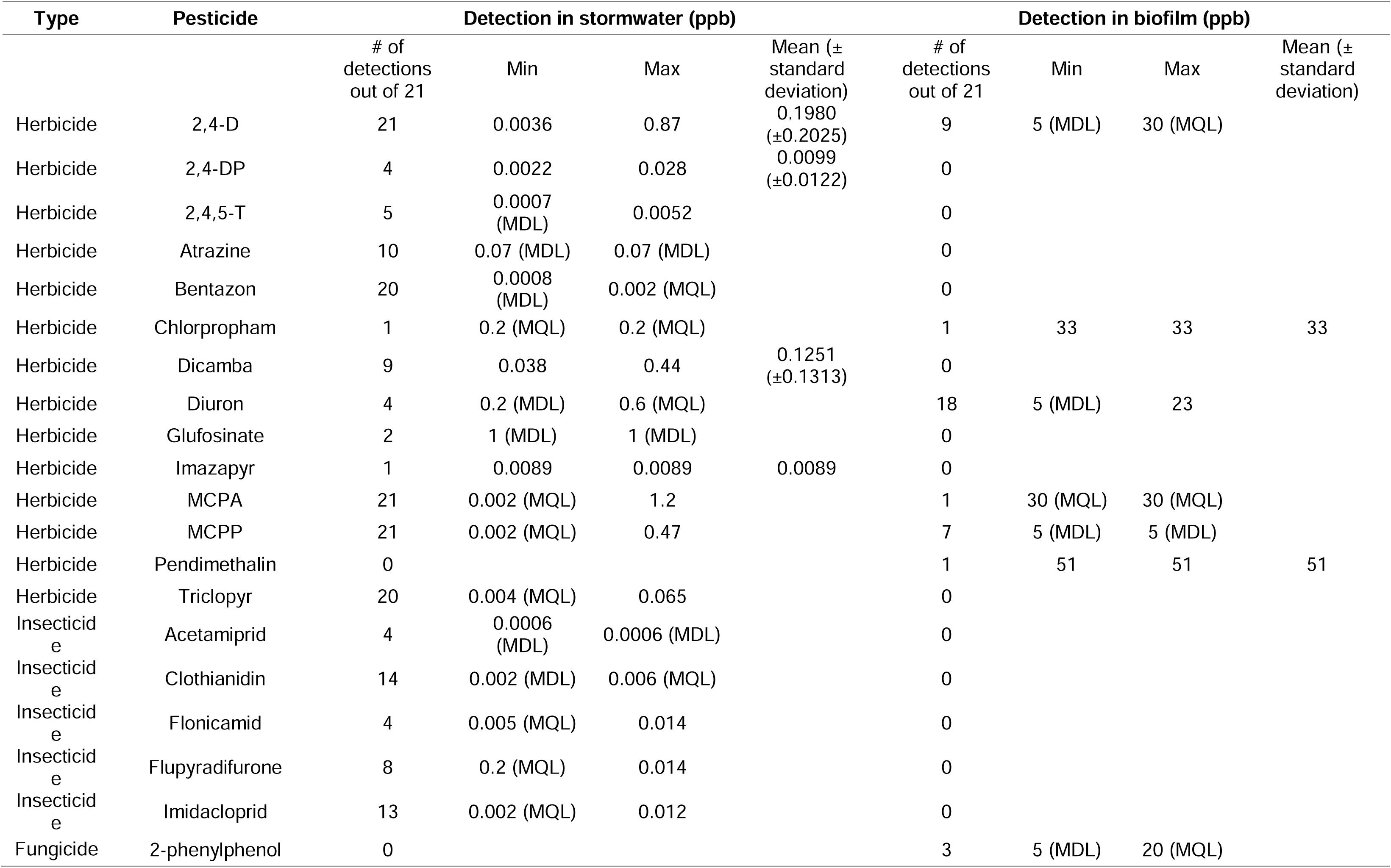
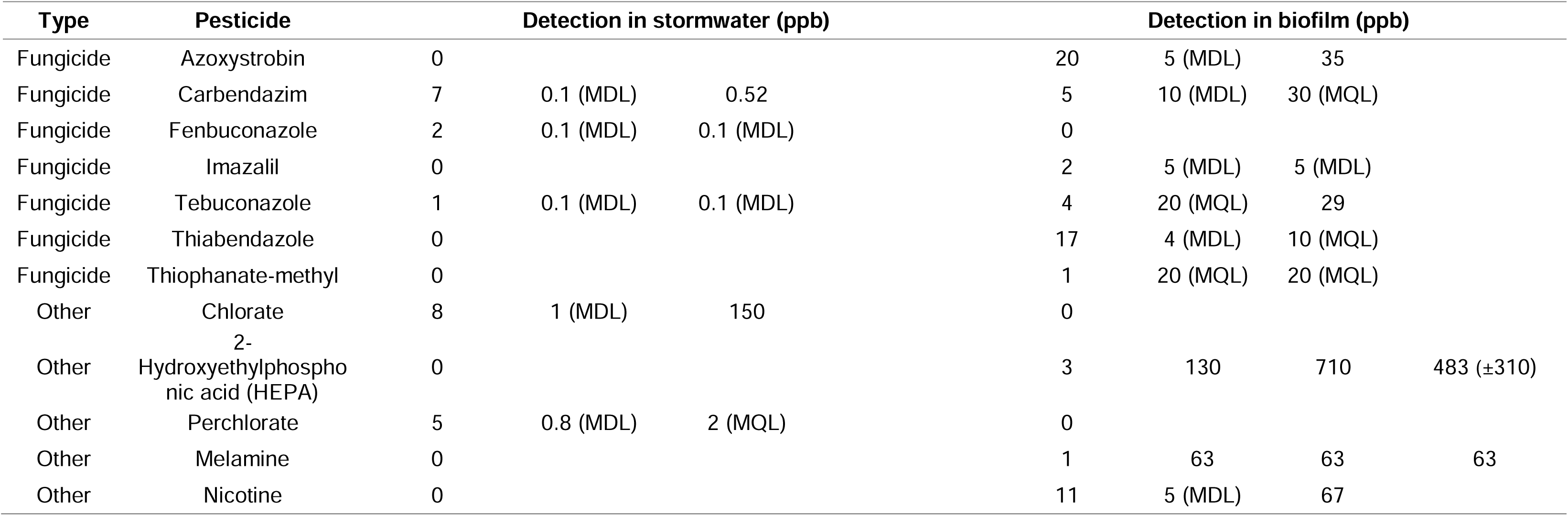
Detection and quantification of 32 pesticides found in water or biofilm samples collected from 21 stormwater ponds in Brampton, Ontario. A full list of the ∼500 pesticides tested for, with their respective limits of detection and quantification, is shown in Appendix A. MDL = Method Detection Limit, MQL = Method Quantification Limit.

Fewer pesticides were detected in biofilm (n = 16) than in stormwater samples (n = 23). This contradicts the findings of Rooney et al. (2020) who detected more pesticides in their biofilm samples (n = 20) than in water samples (n = 9). This contradiction could be explained by the contrasting catchment types, as Rooney et al. (2020) examined an area impacted by agricultural runoff where water moving from fields, through tile drainage, and into ditches, likely accumulates organic matter and transports hydrophobic chemicals that can then adsorb onto biofilms. In the catchments of the Brampton stormwater ponds, stormwater flows rapidly over impervious surfaces like roads, and may be moving primarily water-soluble chemicals that are less likely to adsorb onto biofilms. Application patterns could also play a role: if fewer pesticides with low solubility, for example, are applied in urban areas than in agricultural lands, fewer would be available to be transported into stormwater ponds. Additionally, contrasting limits of detections for the analysis of pesticides in water samples, compared with biofilm samples, may obscure the actual proportion of pesticide occurrences in each matrix type.

Figure 2A and 2B illustrate the distribution of pesticide detections in each matrix by pesticide type. A greater number of herbicides were detected in water (n = 13) than in biofilm (n = 6), while the opposite was true for fungicides (n = 7 in biofilm versus n = 3 in water; Table S2). Five insecticides were detected in water samples, while none were detected in biofilm samples. We identified five other biocides in total, 2 in water samples and 3 in biofilm samples.

**Figure 2.**
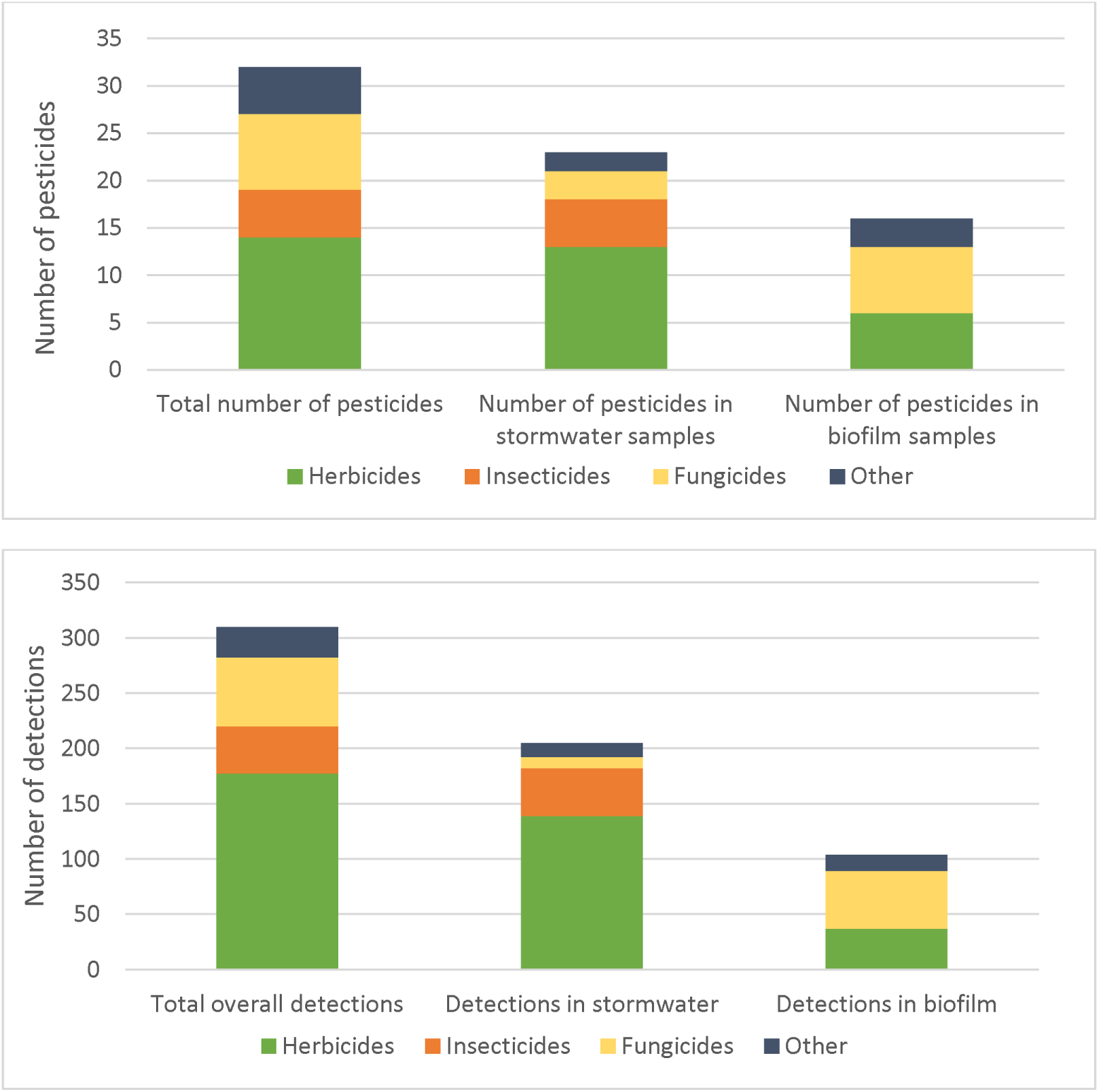
(A) Number of different pesticides detected in stormwater samples (n = 21) and biofilm samples (n = 21). (B) Number of pesticide detections in stormwater and biofilm samples across 21 stormwater pond sites.

Despite differing sampling approaches and suites of pesticide analytes, our herbicide detections were comparable to the only other known publication to report pesticide presence in water samples in Ontario’s urban stormwater ponds. Of the 35 herbicides and insecticides analyzed, Struger et al. (1994) similarly reported a prevalence of phenoxy acid herbicides, but also a collection of pesticides analyzed but not found in our samples (e.g., metolachlor, diazinon, and trifluralin). Pesticide-use profiles have certainly changed since 1994 because of since-introduced legislation (e.g., the use of pesticides for cosmetic purposes was banned in 2009 Ontario under the Pesticides Act; R.S.O. 1990, c. P.11) as well as an increase in public awareness of environmental concerns from pesticide contamination (Arya 2005; Cole et al., 2011). However, despite Ontario’s current cosmetic pesticide-use ban, pesticide application is widespread in urban areas, including in turf management, in construction materials, in pest controls, municipal lands (e.g., right-of-ways), and in other non-cosmetic applications (Pesticides Act, R.S.O. 1990, c. P.11). Additionally, pesticide products used in urban applications may have different formulations but contain the same active ingredients as products restricted to agricultural uses. This could explain the commonalities between the pesticides we detected and those commonly reported in studies based in agriculturally dominated landscapes, even if the urban-use products contain lower concentrations of the active ingredient or different adjuvants than their agricultural counterparts. For example, in an agriculturally impacted coastal marsh, Rooney et al. (2020) found a similar dominance of herbicides in their water samples, compared to our urban samples. However, these two land use types were distinguished by clear differences in the ratios of insecticides and fungicides between water and biofilm samples. This pattern illustrates two contrasting pesticide use and fate profiles that are important for characterizing pesticide contamination across landscapes.

We also grouped pesticides based on their registration status in Canada, shown in Table 3, with most of the pesticides we detected (26 out of 32, or 81%) falling into the current-use category, one historic-use pesticide (2,4,5-T), and the remaining five with either unlisted (imazalil, HEPA, and perchlorate) or non-pesticidal uses (melamine and nicotine). Historic-use pesticides, such as DDT, are no longer permitted for use in the province. Although historic-use pesticides have been reported in contemporary samples of water (Raby et al., 2022) and biofilms (Rooney et al., 2020) collected from agriculturally impacted environments, we did not expect to find many historic-use pesticides in our stormwater ponds. Although Brampton is surrounded to its north and west sides by agricultural lands, the majority of stormwater contributed to stormwater ponds is delivered via the underground stormwater system in its sewer shed and so stormwater entering the ponds has not typically passed over agricultural lands where it could pick up sediments contaminated with historic-use pesticides. The category containing compounds with unlisted or non-pesticidal uses could be associated with chemicals unique to or more prevalent in urban areas; for example, melamine is found as a component of many common household products (e.g., kitchenware, furniture, and cleaning products), as a nitrogen-based flame retardant, and in the manufacturing of plastics (PubChem). Importantly, pesticides in the unlisted or non-pesticidal use categories were detected more frequently in biofilm than in stormwater, necessitating the inclusion of biofilm sampling in urban monitoring to provide comprehensive assessment of contaminants.

**Table 3.**
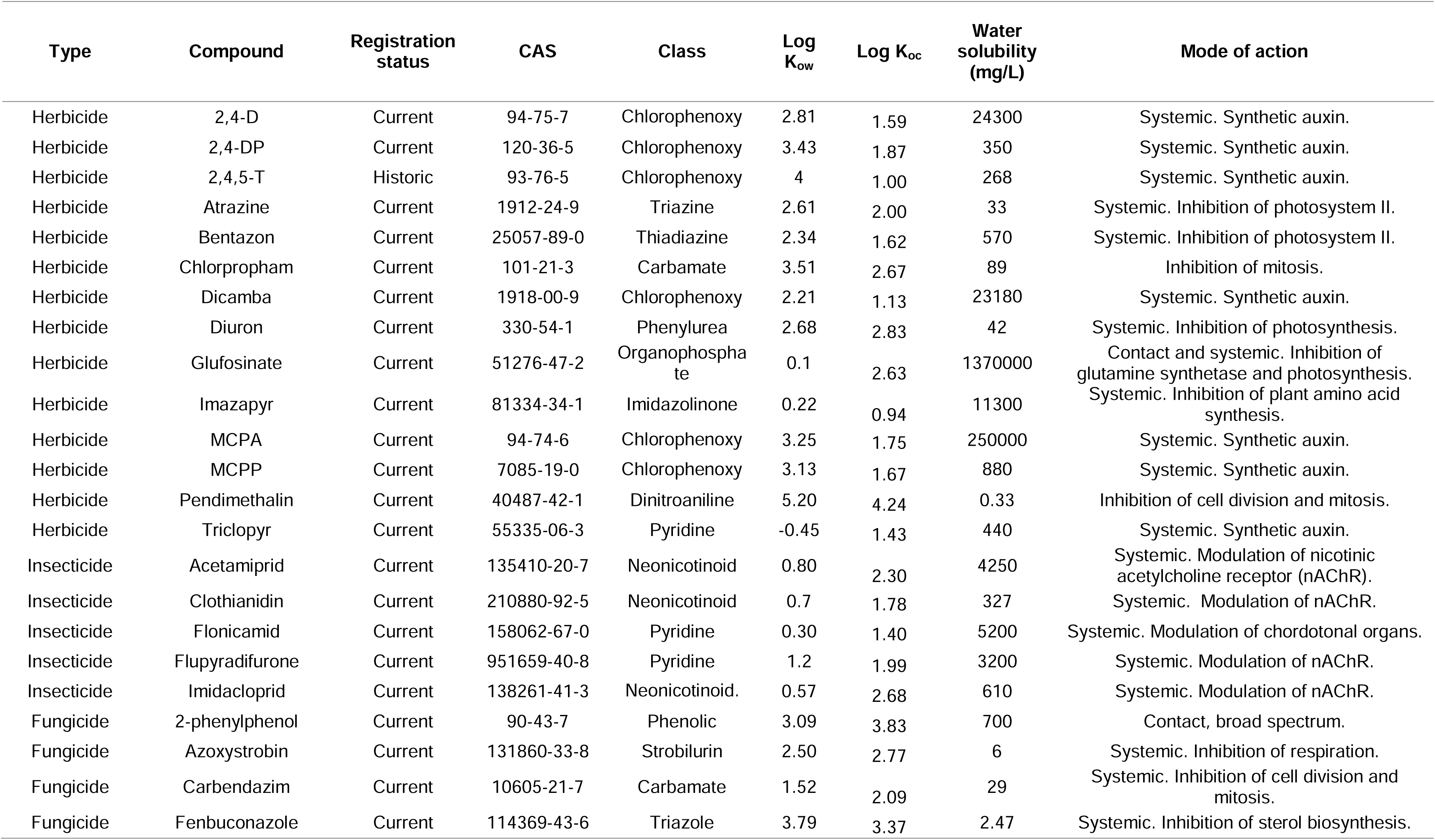
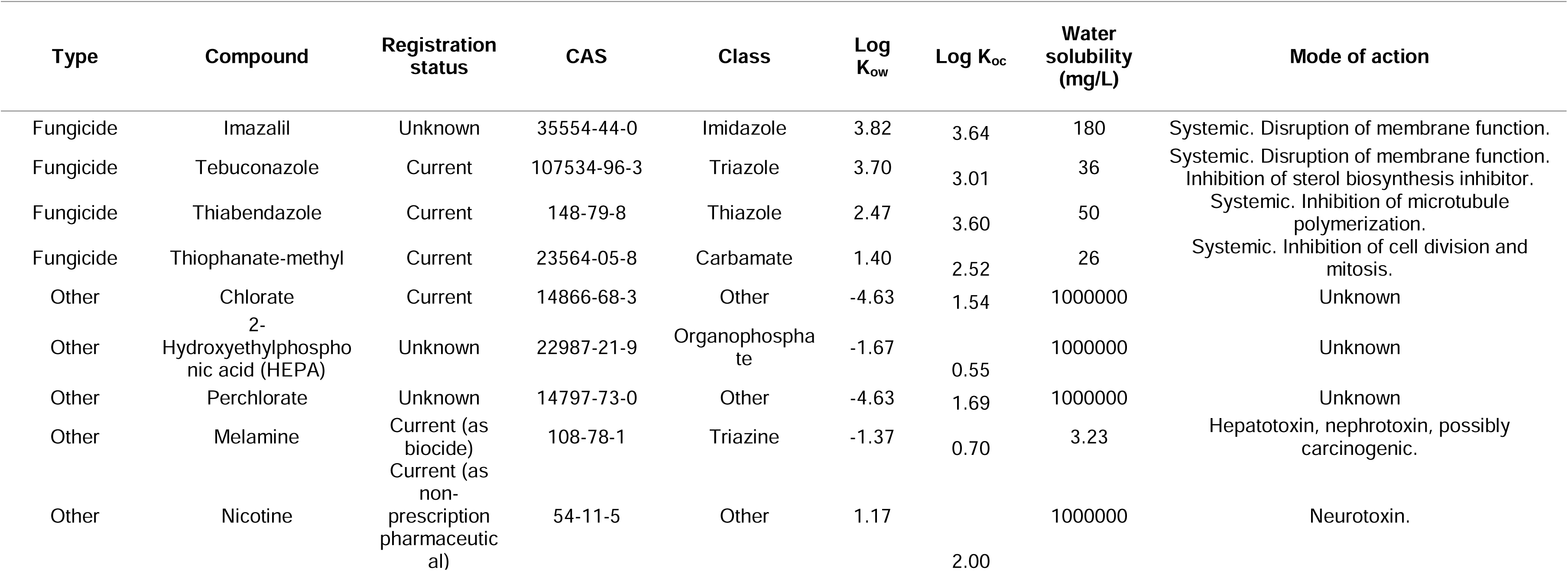
Chemical properties and other notable characteristics of pesticides detected in water and in biofilm. CAS number, chemical class, chemical properties, and mode of action for each pesticide were obtained from the Pesticide Properties Database, PubChem online database, and ChemSpider online database. Current registration status derived from Health Canada’s Pesticide Product Information Database.

#### 3.1.2 Potential risks to biota

Although substantial concern surrounding contaminated stormwater centers on effects to downstream habitats (e.g., Carpenter et al., 2016), stormwater ponds are important hosts for urban biodiversity and should also be subjected to contaminant risk assessments (Rooney et al., 2015; McIsaac 2022; Bradley et al., 2023). While pesticides were ubiquitous in our assessment of stormwater ponds, concentrations were generally low with most water and biofilm samples being below the limits of quantification. However, some pesticides can be toxic at very low levels (e.g., EC50 = 1.6 µg L^-1^ for *Neocloeon triangulifer* exposure to acetamiprid; Raby et al., 2018), thus we assessed whether the pesticides we observed in stormwater ponds could present a threat to aquatic biota.

Of the 16 compounds detected in biofilm samples, 9 (56%) were not detected in stormwater samples. Failure to capture these pesticides through water sampling methods suggests an underestimation of the toxicological burden experienced by aquatic biota and represents a critical gap in our understanding of pesticide contamination in ecosystems. Unfortunately, there remains a lack of available and appropriate dietary toxicity thresholds relevant to aquatic species. As such, it is difficult to estimate the proximity of quantified pesticide levels in biofilm to concentrations causing harm to the growth or survival of its consumers. Testing of dietary consumption of pesticide-contaminated biofilm is a novel ecotoxicological approach (Ijzerman et al., 2023), and thus we are unable to find suitable thresholds to which we can compare our pesticide concentrations and provide an estimation of risk. In future work of this study, we intend to investigate further into the dietary consumption of contaminated biofilm. Hereafter, we discuss the potential risk to biota solely in the context of water exposure.

Many of the detected herbicides are synthetic auxins (see Table 3) meaning they regulate plant growth hormones. For example, 2,4-D causes dysregulation of ethylene production resulting in damaged vascular tissue in broad-leafed weeds such as dandelions (PPDB, 2023). These effects are not seen in narrow-leaf plants like grass, supporting its use in lawncare and turf management, and perhaps explaining their prevalence in stormwater ponds. This specificity also means that 2,4-D, along with most other synthetic auxins, have comparatively lower toxicity to algae: the 96-hour NOEC for *Chlorella vulgaris* is reported as 1 x 10^5^ ppb, whereas the 7-day EC50 for *Lemna gibba* is reported as 2700 ppb (PPDB, 2023). The maximum concentration of 2,4-D found in our samples was 0.87 ppb, and other synthetic auxins similarly fell substantially below reported toxicity thresholds. On the other hand, photosynthesis inhibitors were also observed in our stormwater pond samples. These affect both plants and algae as this mode of action hinders photosynthetic activity in most primary producers. The remaining herbicides we detected included those with modes of action targeting mitosis and amino acid synthesis. Many of these herbicides fell below the MDL or MQL, and all observations were below reported thresholds for the protection of aquatic biota.

As a key part of an insects’ central nervous system, the nicotinic acetylcholine receptor (nAChR) is targeted by many synthetic insecticides, including the majority of those detected in our samples (PPDB, 2023). Modulation of this neurotransmitter receptor is fairly specific to insects but may pose a risk to other aquatic organisms. The presence of neonicotinoids in stormwater is not surprising, given previous detections in Ontario urban and rural surface waters (Metcalfe et al., 2019; Raby et al., 2022), however measured concentrations were well below known toxicity thresholds. For example, the highest concentration of flonicamid was 0.014 ppb, which was considerably lower than the chronic NOEC for *Daphnia magna*, 3100 ppb (PPDB), and the USEPA Aquatic Life Benchmark (ALB) for chronic exposure to freshwater invertebrates, 3000 ppb (USEPA). Imidacloprid, measured in our stormwater samples at 0.012 ppb, was below reported toxicity thresholds: the acute LC50 for *Chironomus riparius* is 55 ppb (PPDB), and the CCME guideline for the protection of aquatic life is 0.23 ppb (CCME). However, this concentration was slightly higher than the USEPA ALB for chronic exposure to freshwater invertebrates, at 0.01 ppb. This was the only measured contaminant in stormwater samples to surpass a benchmark.

Carbendazim was the only fungicide that we detected above its quantification limit in stormwater samples. The maximum concentration we observed was 0.52 ppb. This is higher than reported by Metcalfe et al. (2016) in their surveys of urban surface waters (0.18 ppb) and yet it remains below the most sensitive threshold for aquatic biota, reported as 1.5 ppb for *Daphnia magna* (NOEC; PPDB). Other biocides detected in stormwater also fell below USEPA benchmarks and PPDB reported thresholds.

Concentrations of pesticides in the stormwater samples were low, never exceeding thresholds or guidelines for ecotoxicological risk to aquatic biota reported by the PPDB and CCME, with only one exceedance of a USEPA benchmark. However, these contaminants were always present as mixtures: a minimum of four pesticides were detected at each pond, and 57% of ponds had 10 or more pesticides detected (See SI for pesticide detections and concentrations in water samples). The mixtures included multiple pesticide types (herbicides, fungicides, insecticides) and multiple modes of action within a single type (e.g., synthetic auxins and photosynthesis inhibitors). Such complex mixtures may have greater toxicity to biota than predicted from any single pesticide as their toxicities may combine additively or synergistically (Hernández et al., 2013; Weisner et al., 2021). Consequently, despite the low concentrations detected in our samples, the ubiquity of pesticides and the diversity of mixtures detected in our study raises concern for aquatic biota in stormwater ponds and emphasizes the need for more research on pesticide mixtures (Hernández et al., 2017).

### 3.2 Relationship between occurrences of pesticides in each matrix and their physicochemical properties

We evaluated whether physicochemical properties of pesticides that are commonly invoked to understand their fate in aquatic environments (Table 3) were predictive of the number of detections of pesticides in stormwater and in biofilm from stormwater ponds using generalized linear models (Table 4). We found that water solubility did predict the number of pesticides detected in water samples (Figure S2B), but not in biofilm samples (S2A). The lack of relationship between water solubility and detection in biofilms could be due to the heterogeneous surface of biofilm, which provides diverse chemical sorption sites that contribute to its ability to accumulate a wide assortment of contaminants (Bonnineau et al., 2021). For example, although pesticides with lower water solubilities were indeed represented in biofilm samples, HEPA and nicotine, the two most soluble compounds detected across all sites with water solubilities over 1 x 10^6^ mg L^-1^, appeared exclusively in biofilm samples.

**Table 4.**
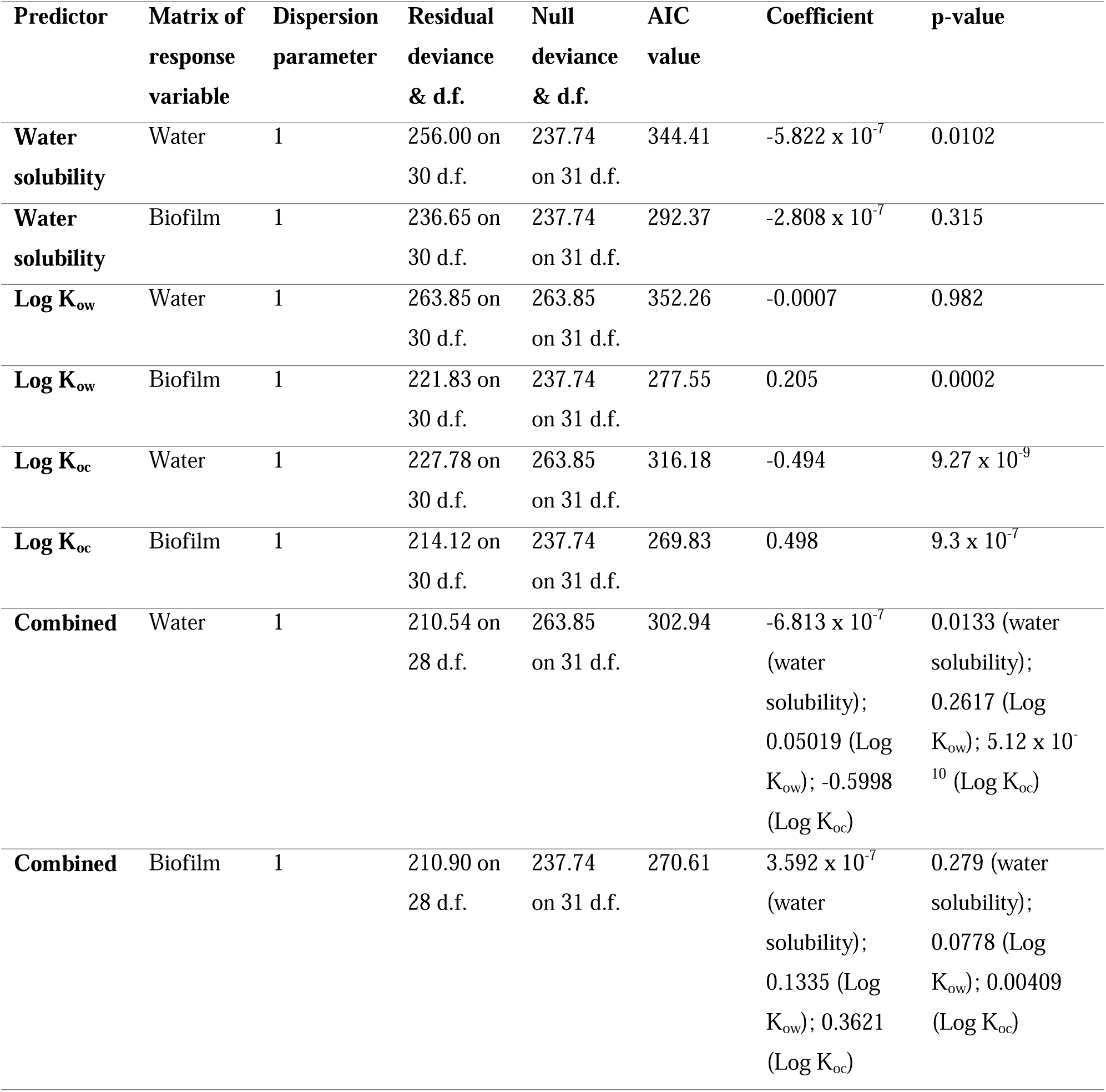
Results of generalized linear models with gamma distributions for pesticide physicochemical properties with detections in water and in biofilm.

The octanol-water partition coefficient (K_ow_) indicates the ratio of a chemical’s concentration in the non-polar octanol phase to its concentration in the polar aqueous phase of an octanol-water two phase system. Log K_ow_ was predictive of the number of pesticides detected in biofilm (Figure S2C), but not in water (Figure S2D). The Log K_ow_ is a metric of hydrophobicity and is often used to calculate the accumulation of a chemical into living biological material as a bioconcentration factor (e.g., Mackay & Fraser, 2000) Thus, we expected that pesticides with higher Log K_ow_ would more readily be accumulated out of the water phase and onto or into the living biofilms.

The soil adsorption coefficient (K_oc_) is the ratio of a chemical’s concentration in a soil or sediment phase to its concentration in a water phase. In our context, this coefficient represents the adsorption of pesticides onto the organic matter contained within biofilms, both intra– and extracellularly. We found that Log K_oc_ predicted the number of detections in both biofilm (Figure 3A) and water (Figure S2E). We expected that pesticides with higher Log K_oc_ values may bind to organic-rich materials more strongly and thus have elevated occurrences in biofilms, while pesticides with lower Log K_oc_ values and weaker bonds with organics should be more dissolved and mobile in the water.

**Figure 3.**
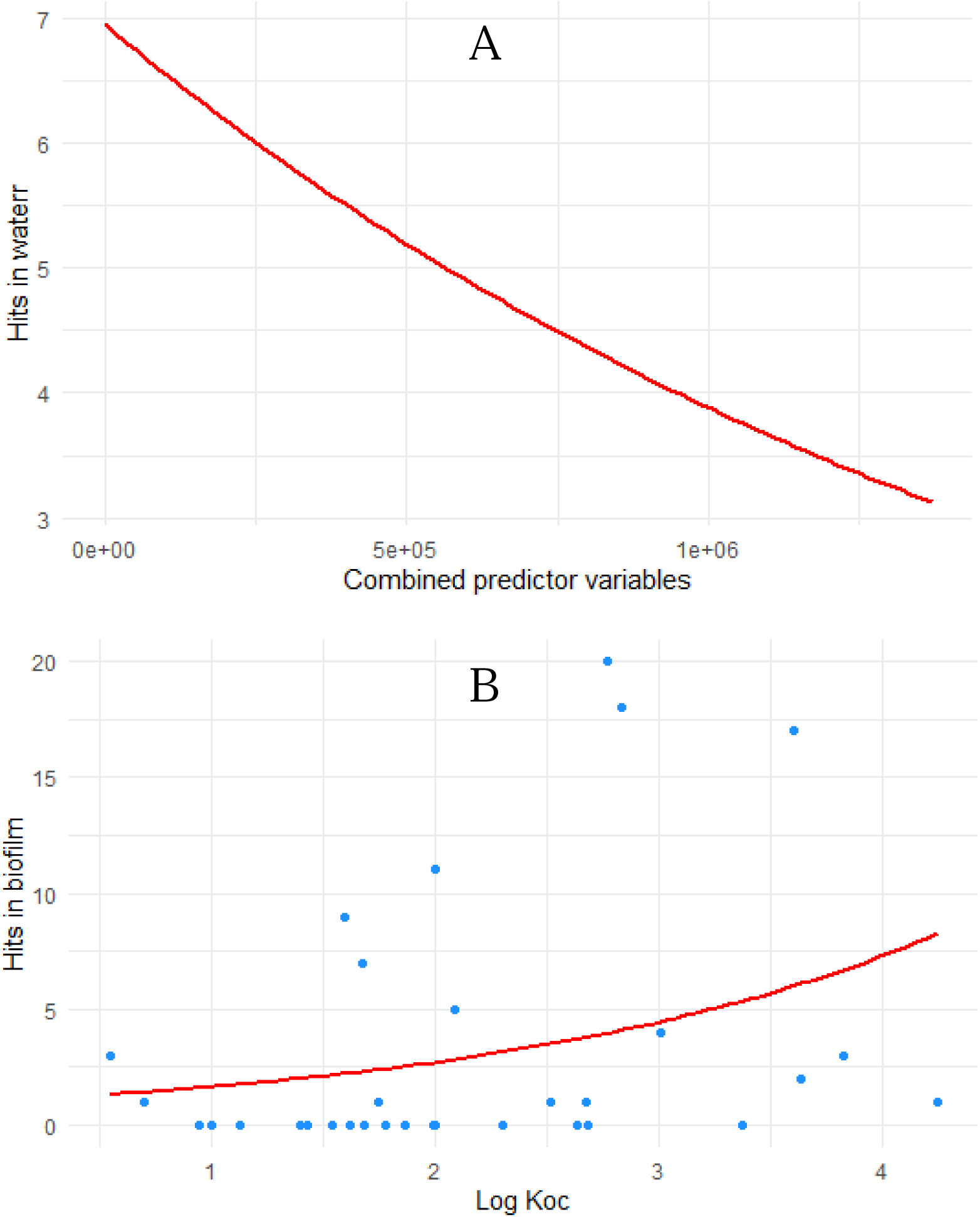
Generalized linear models that best predict the number of pesticides detected in water and biofilm samples. A combined model using water solubility, Log K_ow_ and Log K_oc_ best predicted the number of pesticides detected in water (A), while of the number of pesticide detections in biofilm was best predicted by Log K_oc_ (B).

Additionally, we tested combined models using all three variables as predictors of pesticide occurrences in stormwater (Figure 3B) and in biofilm (Figure 2SF). The model for detections in water and the model for detections in biofilm samples had delta-AIC values of 13.24 and 0.78, respectively (raw AIC values shown in Table 4). This indicates that the number of pesticide detections in water is better predicted by a combined model than by individual pesticide properties, but that the number of pesticides detected in biofilm is not better predicted by the combination of three pesticide properties than it was by Log K_oc_ alone.

### 3.3 Bioconcentration factors

#### 3.3.1 Calculated and estimated bioconcentration factors

All BCF values, both calculated and estimated, as well as averages for calculated BCFs and residuals for the pesticides with multiple calculated BCFs, are reported in Table 5. Calculated BCFs spanned a wide range (4.2 – 1275). Of the 42 calculated and estimated BCFs, 10 were attributed to herbicides, 18 to fungicides, and 14 to other biocides. No insecticides were found to have bioconcentrated in the biofilm. Bioconcentration factors (BCFs) are valuable indices for comparing relative concentrations of pesticides in two matrices and for quantifying accumulation from ambient water. Recent studies have quantified the magnitudes of pesticide accumulation in biofilm (e.g., Mahler et al., 2020; Rheinheimer dos Santos et al., 2020), however only two other studes explored a similarly large set of pesticides and calculated bioconcentration factors (BCFs; Rooney et al., 2020; Ijzerman et al., 2023). Our study is the first to document pesticide BCFs in biofilm in a non-agricultural context.

**Table 5.**
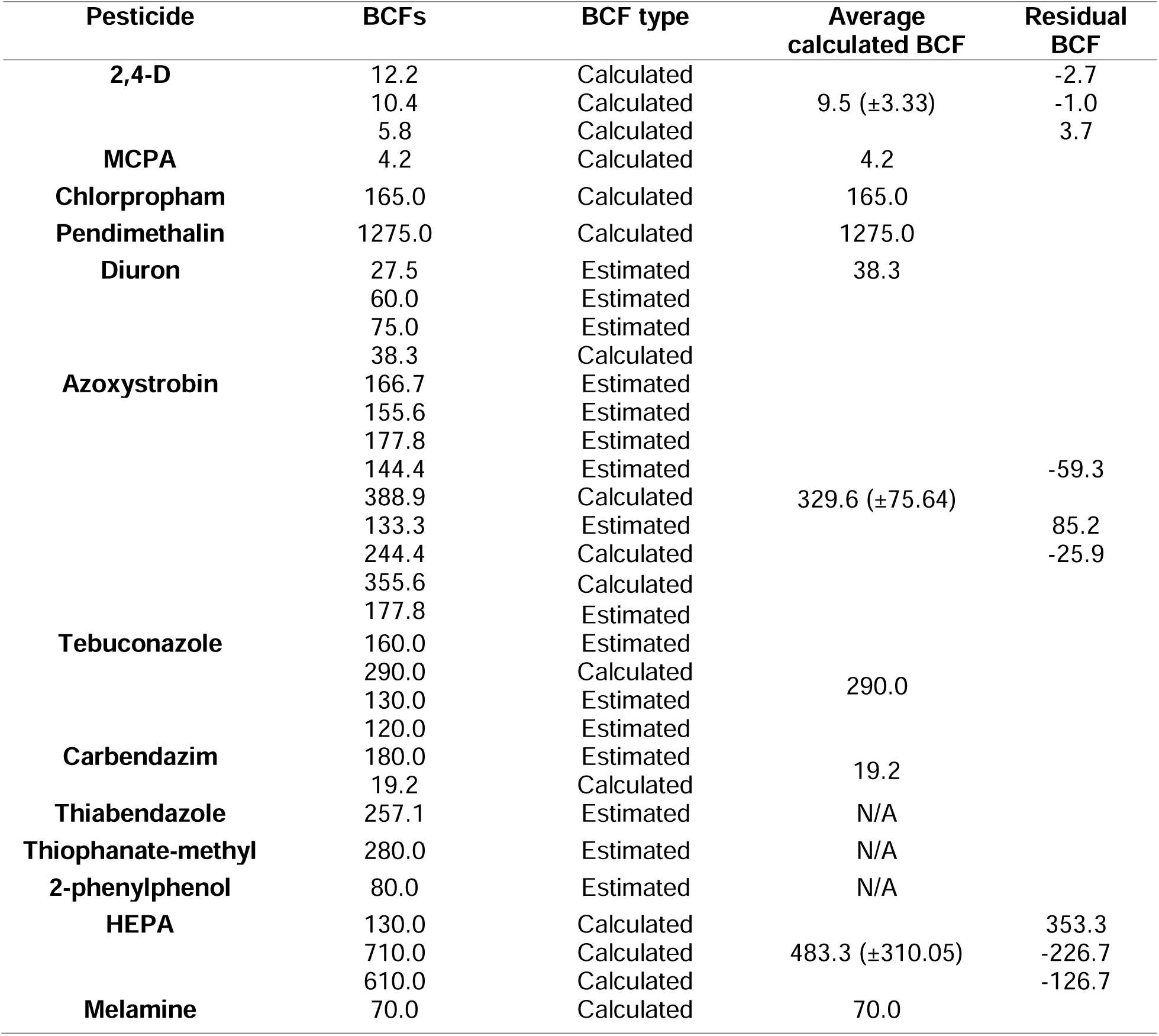
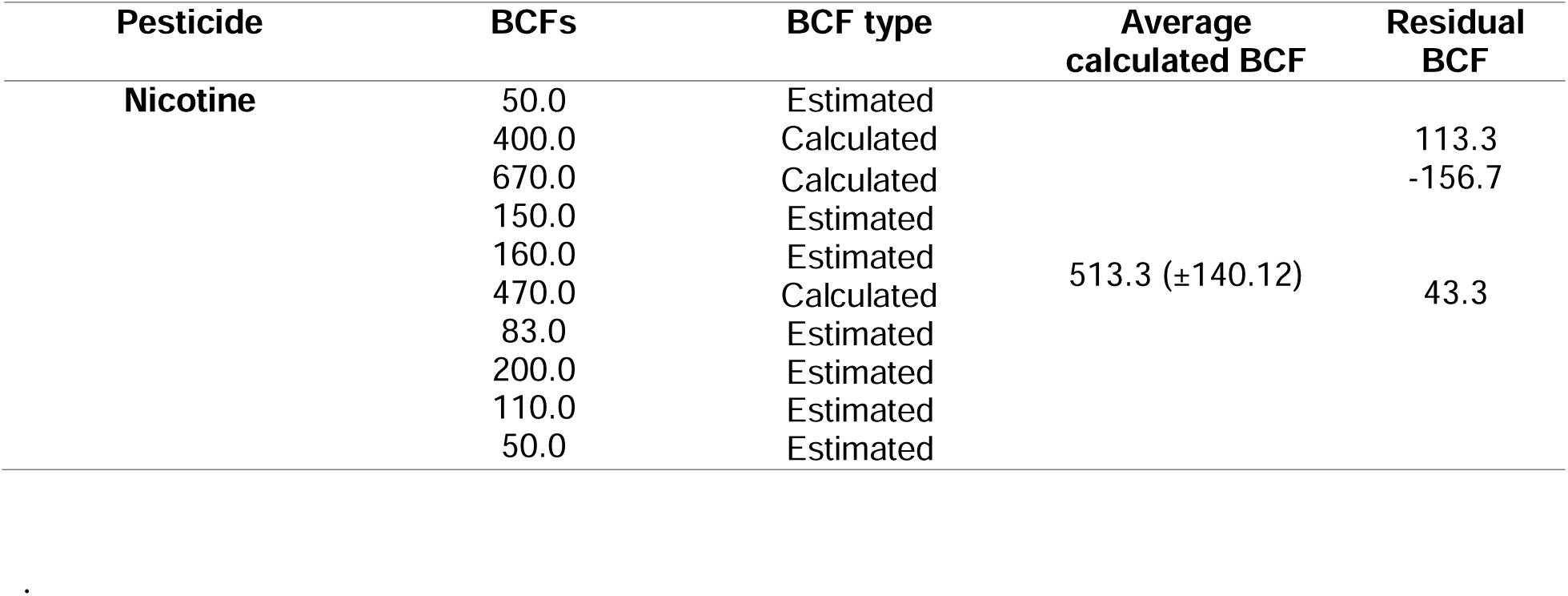
Calculated and estimated bioconcentration factors (BCFs) of pesticides quantified in stormwater ponds.

The BCF values reported in pesticide databases (such as the PPDB) for pesticides in aquatic ecosystems are typically based on fish. Although pesticide physicochemical properties are thought to dictate pesticide accumulation in aquatic organisms (Mackay & Fraser 2000; Katagi 2010), the diversity in characteristics of the biofilm surface and EPS matrix that orchestrate sorption and binding of pesticides (Lundqvist et al., 2012) could result in differences from fish BCFs. However, these indices are still useful for contextualizing our reported BCF values. While a few differences exist, we observed surprising agreement in the order of magnitudes of our reported BCF values and those reported by pesticide databases and by other scientific studies (e.g., Rooney et al. 2020). Given the variability in exposure levels, water quality, and the composition of biological tissues being considered by these information sources, we propose that only differences in the order of magnitude of BCF values are meaningful.

Our results generally agreed with those of Rooney et al. (2020), who also found that biofilm accumulates a wide range of pesticides from the water of coastal marshes, however our reported BCF values were more consistent with those recorded for pesticides in the PPDB. For example, for 2,4-D, the PPDB lists a BCF in fish of 10, consistent with our average BCF of 9.5 (± 3.33), yet lower than reported by Rooney et al. (2020), who calculated a BCF of 42 in freshwater periphyton. Similarly, our lowest recorded BCF, 4.2 for MCPA, was comparable to the PPDB reporting of 1 in fish, but again differs substantially from the BCF of 2459.2 reported by Rooney et al. (2020). Atrazine was detected in 48% of stormwater samples but did not accumulate in our biofilm samples, also contradicting accumulation in periphyton found by Rooney et al. (2020; BCF = 112 – 350) and Nikkila et al. (2001; BCF = 215 – 350). These differences could be driven by environmental factors, such as catchment type or water quality, however more investigation is needed to understand the mechanisms of pesticide accumulation before we can determine the influence of external conditions.

No literature was found to measure the BCF of chlorpropham, but the PPDB listing for fish (144) was comparable to our calculation (165). A geometric mean BCF of 1878 for pendimethalin in aquatic organisms determined by Vighi et al. (2017) is likely a more appropriate comparison point than the BCF reported for fish in the PPDB, 5100, although both are within the same order of magnitude as our calculation of 1275. Diuron was measured to be bioconcentrated at a factor of 2 in freshwater fish by Call et al. (1987) and is reported by the PPDB as 9.45 in fish, both lower than our calculated BCF of 38.3.

Similarly, BCFs for fungicides reported by the PPDB and most accessible literature (e.g., Andreu□Sánchez et al., 2012; Ju et al., 2019) measured accumulation in either fish or crop plants, the exception being those reported in periphyton by Rooney et al. (2020). Calculated BCFs for azoxystrobin and tebuconazole (688.0 and 712.9, respectively) by these researchers were higher yet within the same order of magnitude as our BCF calculations (329.6 and 290.0, respectively). This discrepancy (as with herbicidal BCFs) is possibly due to higher loads of agricultural pesticides delivered to their sampling environment. This explanation is substantiated by Ding et al. (2019), who described a wide range of cellular accumulation of carbendazim (BCFs = 12.1 – 1336.2) in dried algal samples when exposed to a range of concentrations over the span of 144 hours. The relationship between cellular accumulation of carbendazim and initial exposure concentrations in Ding et al. (2019) may explain variations in periphyton or biofilm accumulation in environments with different exposures to pesticide loads. Both our calculated carbendazim BCF (19.2), and that reported by the PPDB (25), fell into the range given by Ding et al. (2019). Thiabendazole and thiophanate-methyl had estimated BCFs (257.1 and 280.0, respectively) higher than those reported by the PPDB for fish (96.5 and 75, respectively). No relevant BCFs for the remaining biocides were found in literature, although the PPDB reports a BCF of 21.7 (for an unknown organism) for 2-phenylphenol, which we estimated at 80.0 in our biofilm samples.

### 3.4 Chemical and environmental drivers of bioconcentration factors

#### 3.4.1 Bioconcentration factors and pesticide physicochemical properties

We used generalized linear models to investigate how pesticide physicochemical properties influenced the magnitude of pesticide bioconcentration, the results of which are presented in Table 6. We did not find calculated BCF values to be significantly predicted by water solubility (Figure S3A), Log K_ow_ (Figure S3B), or Log K_oc_ (Figure S3C) alone. A model combining these three properties yielded a better fit (delta AIC = 3.74; Figure 4) than any of the three properties modelled alone, but within this best fitting model, only water solubility had a p-value below 0.05 (Table 6). This reveals that the three physicochemical properties were not good predictors of pesticide BCFs in stormwater pond biofilms.

**Figure 4.**
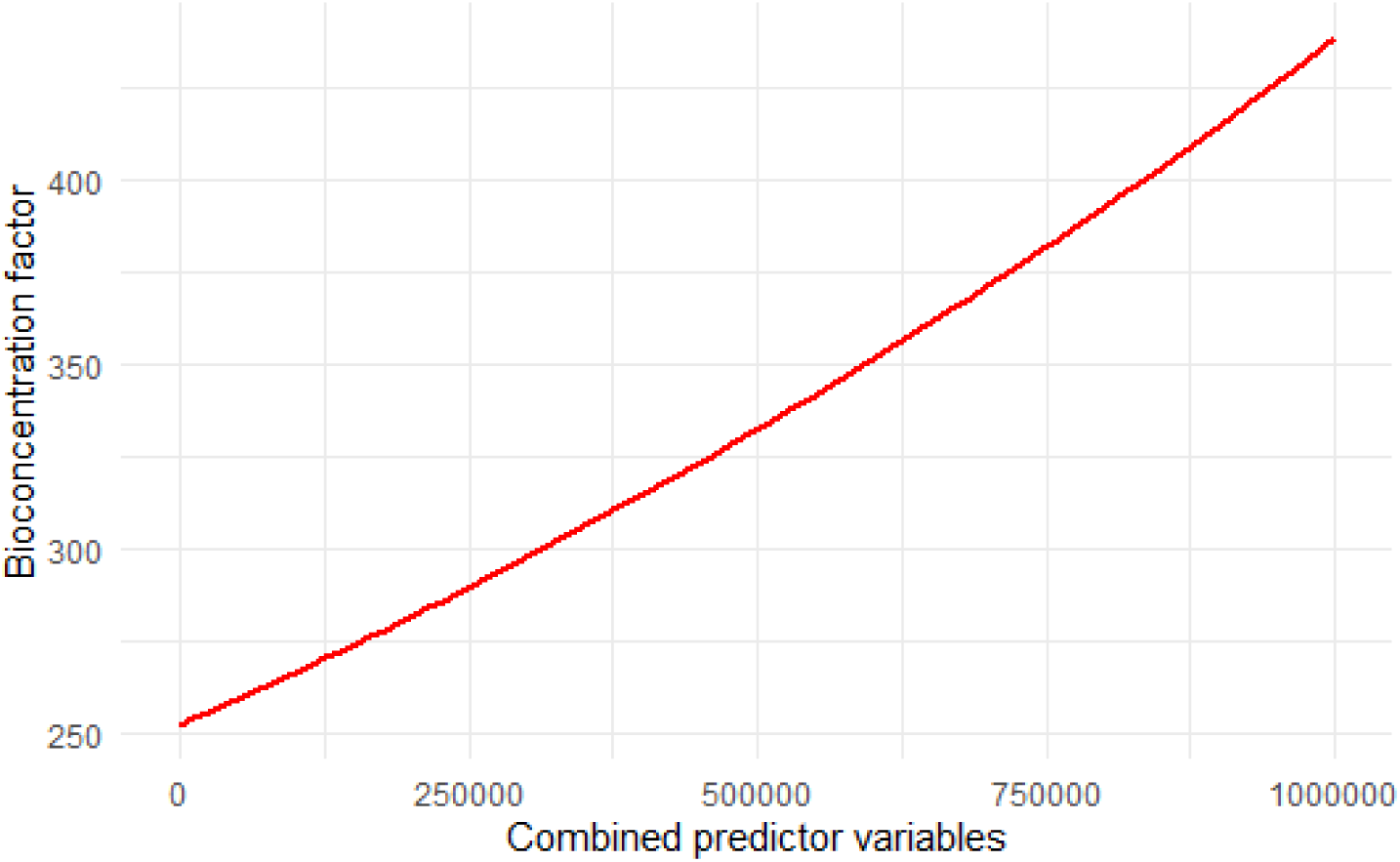
A combined generalized linear model using water solubility, Log K_ow_ and Log K_oc_ as predictors of the bioconcentration factor of pesticides.

**Table 6.**
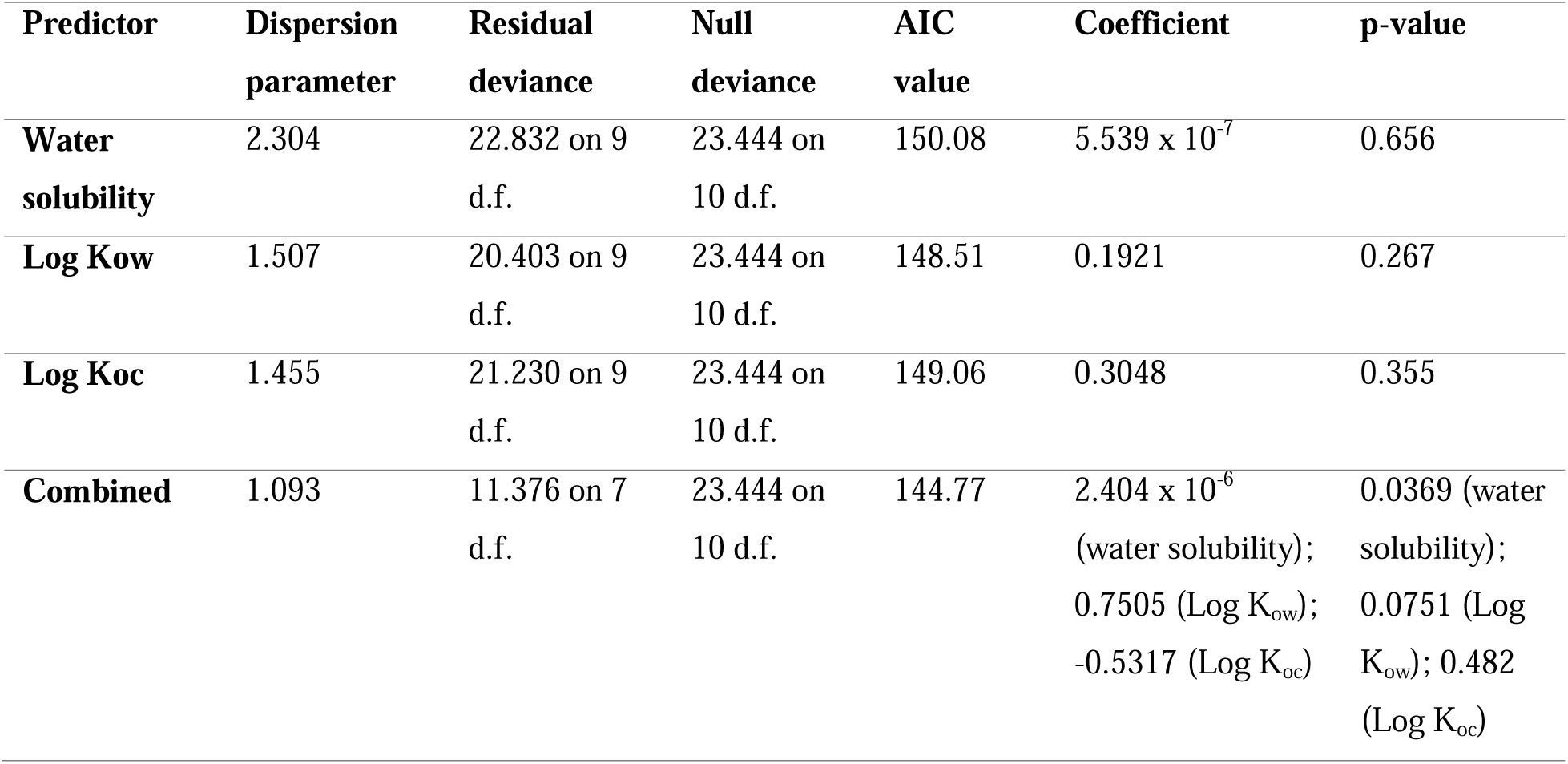
Results of generalized linear models for analysis of pesticide physicochemical properties as predictors of bioconcentration factors. Asterix indicates statistical significance (p < 0.05).

Although Lundqvist et al. (2012) and Nikkila et al. (2001) found hydrophobicity was related to pesticide accumulation, Rooney et al. (2020) did not discern any strong physicochemical drivers of pesticide BCFs in periphyton. Our generalized linear regressions (Figure S3) agree with the findings of Rooney et al. (2020). From this, we conclude that BCF values for pesticides accumulating in biofilms cannot reliably be estimated from physicochemical properties and that validation of pesticide BCFs from field-based biofilm monitoring are required. Biofilm sampling is thus an important addition to contaminant monitoring.

#### 3.4.2. Bioconcentration factors and other parameters

As pesticide properties were not the main drivers of pesticide accumulation in our biofilm samples, we explored whether characteristics of the environment might better predict BCF variance across the stormwater ponds. The results of the generalized linear models are presented in Table 7.

**Table 7.**
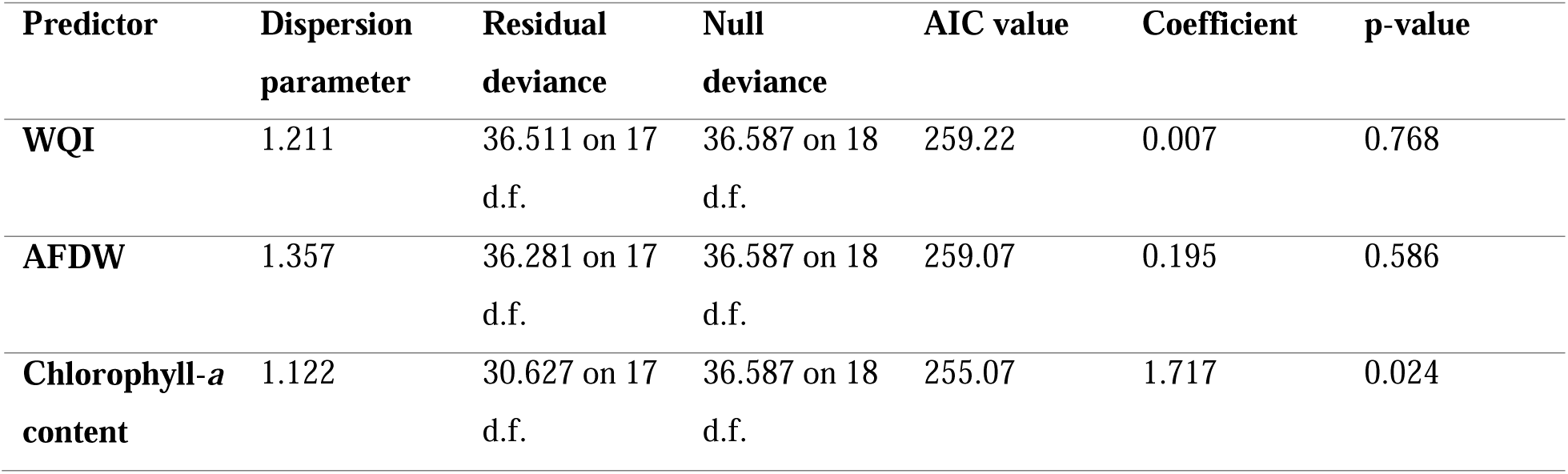
Results of generalized linear models with gamma distributions with water quality index (WQI), ash-free dry weight (AFDW) of biofilm, and chlorophyll-*a* content of biofilm as predictors of calculated bioconcentration factors (BCF).

We did not find a significant relationship between calculated BCF values, and the Water Quality Index (WQI) scores calculated for each pond site (Figure S4). Some BCFs varied widely, such as those calculated for HEPA, which ranged from 130 to 710. We had predicted that these observed variations could be features of the surrounding water quality and its impact on biofilm function and growth rate, resulting in more or less bioconcentration than expected. Deteriorated water quality could shift community composition (Tien et al., 2013) and pesticide tolerances (Larras et al., 2013), potentially leading to changes in accumulation capacities, and factors not included in the WQI score, such as artificial light pollution, could also alter community composition.

We also did not identify a significant relationship between BCFs and ash-free dry weight (AFDW; Figure S5), suggesting there was not a dilution effect (i.e., lower pesticide concentration with higher biofilm mass). We also used ash-free dry weight as a proxy for biofilm growth rate since all biofilm samplers had equivalent incubation periods at each site, and therefore biofilm samplers with higher ash-free dry weights must have grown at a faster rate. If there was a constant rate of pesticide uptake into biofilms, we would expect to see a negative linear relationship between BCF values and ash-free dry weight. The lack of such a relationship implies that the rate of pesticide accumulation is variable but unrelated to the rate of biofilm growth.

We did find a statistically significant relationship between calculated BCF values and the chlorophyll-*a* concentration measured in the biofilm, shown in Figure 5. A greater proportion of fungicides (n = 18) were accumulated relative to herbicides (n = 10), and at higher magnitudes (Table 5). This pattern could explain a tendency towards autotrophy and thus higher content of photosynthetic compounds in biofilms with higher BCFs and could suggest that the community composition of a biofilm influences the capacity for pesticide accumulation. However, an in-depth assessment of microbial diversity is required to understand this relationship, and as such, is recommended here only as an avenue for future research.

**Figure 5.**
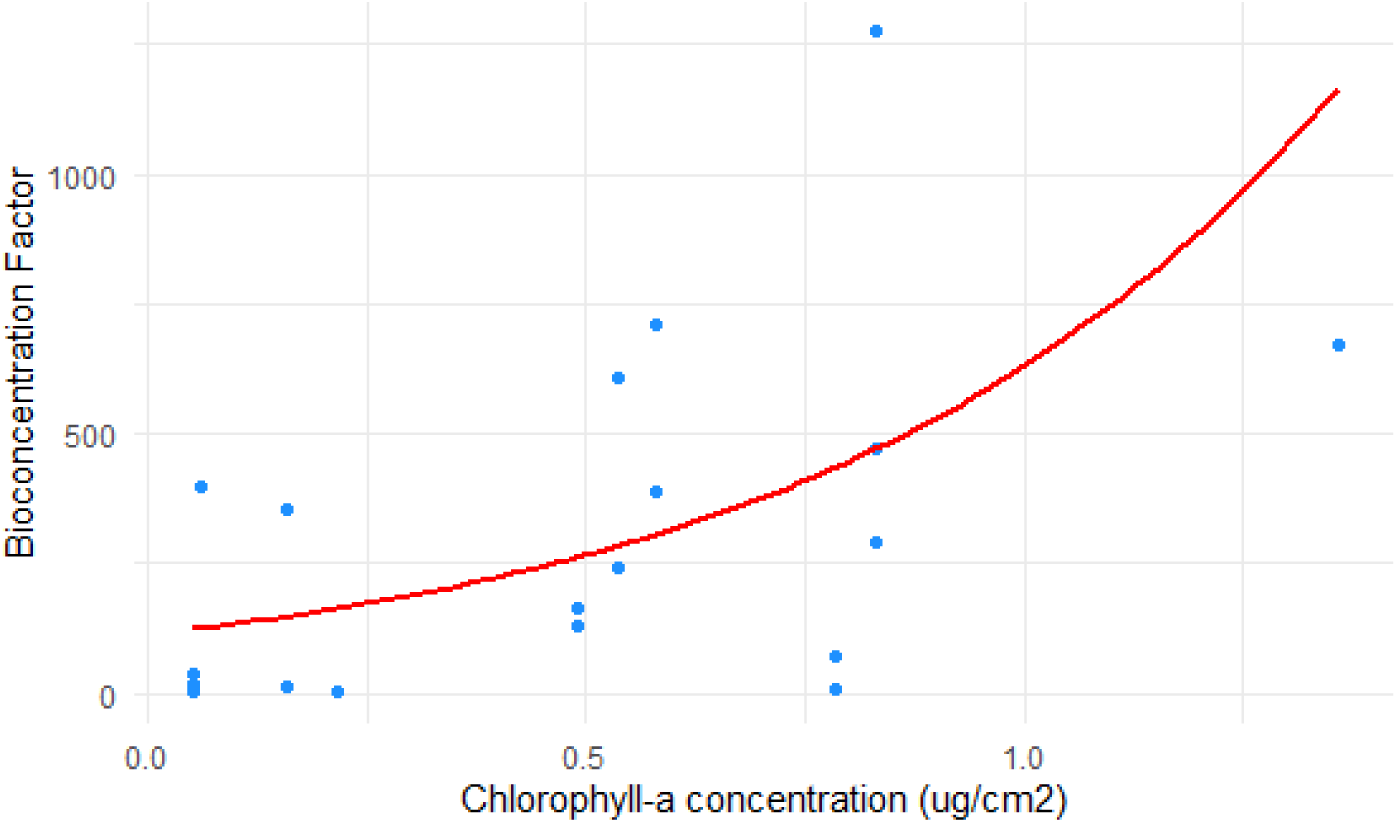
Chlorophyll-*a* concentration (µg cm^-2^) in biofilm and calculated pesticide BCF values (n = 17; p = 0.024) in a generalized linear model.

Our inability to find the expected drivers of accumulation as good predictors of pesticide BCFs is likely due to an insufficient understanding of the mechanisms behind pesticide accumulation in biofilm and thus an incorrect selection of predictor variables. This is still an underrepresented area of research, frequently focusing on a small number of compounds in a controlled environment (e.g., Lundqvist et al., 2012; Chaumet et al., 2019a) and is often limited by technical constraints (e.g., in the quantification of contaminants within the EPS vs intracellularly; Bonnineau et al., 2021). The duration of pesticide exposure (Bonnineau et al., 2021), repartition dynamics between different environmental matrices (Headley et al., 1998), chemical transformation within the biofilms (Carles et al., 2019), and species composition (Lawrence et al., 2001; Berglund et al., 2005) are examples of plausible interacting drivers of pesticide accumulation as identified by other researchers. Yet, disentangling the vast assortment of possible mechanisms driving our calculated BCFs was not feasible in our field-based experiment.

Manipulating the conditions in which a biofilm grows and measuring the responding levels of accumulation of a single pesticide could provide valuable insight into the environmental influences of contaminant-biofilm interactions. For example, in the laboratory, Beecraft & Rooney (2021) found that the BCF of glyphosate in biofilm was dependent on the concentration added, with higher BCFs in lower concentrations, suggesting a saturation effect. This was not a relationship we were able to explore because only one pesticide (2,4-D) had both quantifiable concentrations in water samples and multiple calculated BCF values. Another caveat of our study was our small sample size (n = 17) as we only conducted analyses of possible relationships using our calculated BCF values (not estimated). Future research conducting analyses on a much larger data set could help to better assess relationships between BCFs and pesticide chemical properties, biofilm characteristics, and surrounding water quality in the real world.

## 4. Conclusion and implications for future work

Our study is the first in over 25 years to identify and quantify pesticide contamination in stormwater ponds in Ontario, and the first since key pieces of legislation were put in place to reduce the use of neonicotinoids (OMECP, 2015) and the cosmetic use of pesticides (OMECP, 2014). Although known toxicity thresholds for pesticide exposure in water were not exceeded by levels found in the stormwater ponds, the risks to biota from exposure to complex mixtures and to contaminated biofilm are unknown. The implications of the observed diversity in modes of action for organisms living in these environments is an important area for future research. Our study is also one of the few to report pesticide bioconcentration in biofilm collected from the field. Pesticides, particularly current-use fungicides, were common in biofilms and several pesticides that were present in biofilms were not detected in the water samples, revealing that a comprehensive assessment of pesticide contamination in stormwater ponds really requires biofilm monitoring. The distribution of pesticides in water and biofilm was related to select physicochemical properties, with the Log K_oc_ being a strong predictor in both matrices. Our calculated BCFs indicate pesticide accumulation can vary and may not be predicted by pesticide physicochemical properties or environmental factors, highlighting the need for more research on accumulation mechanisms. The use of biofilm sampling in contaminant monitoring can clearly provide more sensitivity than water sampling in pesticide monitoring programs, especially for chemicals shown to have high BCF values. Water sampling may overlook pesticide risks, as even when pesticides are below detection limits in water samples, biota can still be exposed via consumption of biofilms. This represents an underexplored exposure pathway, and we urge other researchers to consider the potential risks of chronic dietary consumption of pesticides when evaluating toxicological risks and setting ecological thresholds. Additionally, stormwater ponds could be mobilizing contaminants into other urban food webs, as many pond dwellers such as benthic macroinvertebrates and birds act as bridges between aquatic and terrestrial ecosystems. Future work incorporating ecological assessments of these communities could help illustrate the breadth of stormwater contamination.

## Data availability

Supplemental information is provided here: Izma_BCFpaper_supplementalinfo.docx

The data from the present study have been made available on FigShare: https://figshare.com/account/home#/projects/179421 (this isn’t published yet so you may not be able to access it, but links to each dataset are included in the supplemental information document which should be viewable – please let me know if not).

## Acknowledgements

The stormwater pond sites surveyed for this project are situated on the traditional territory of the Anishinaabeg, Haudenosaunee, and Huron-Wendat peoples, where Indigenous communities, including most recently the Mississaugas of the Credit, have lived for thousands of years. The remaining research was conducted at the University of Waterloo, situated on the Haldimand Tract, the land promised to the Six Nations of the Grand River. This land is the traditional territory of the Neutral, Anishinaabeg, and Haudenosaunee peoples, and is now home to many First Nations, Inuit, and Métis peoples.

We thank Brian Atkinson from AFL for conducting all pesticide analyses, Danny McIsaac and Moira Ijzerman for help with site surveying and sample collections, and Hannah Thibault for feedback on an early draft. Biofilm samplers were designed and constructed with the guidance of Hiruy Haile from the University of Waterloo Science Technical Services.

## Author contributions

Gab Izma – Conceptualization, Investigation, Methodology, Formal Analysis, Writing – Original Draft Melanie Raby – Conceptualization, Resources, Funding Acquisition, Methodology, Supervision, Writing – Review & Editing

Ryan Prosser – Conceptualization, Resources, Funding Acquisition, Methodology, Supervision, Writing – Review & Editing

Rebecca Rooney – Conceptualization, Resources, Funding Acquisition, Methodology, Project Administration, Supervision, Writing – Review & Editing

---

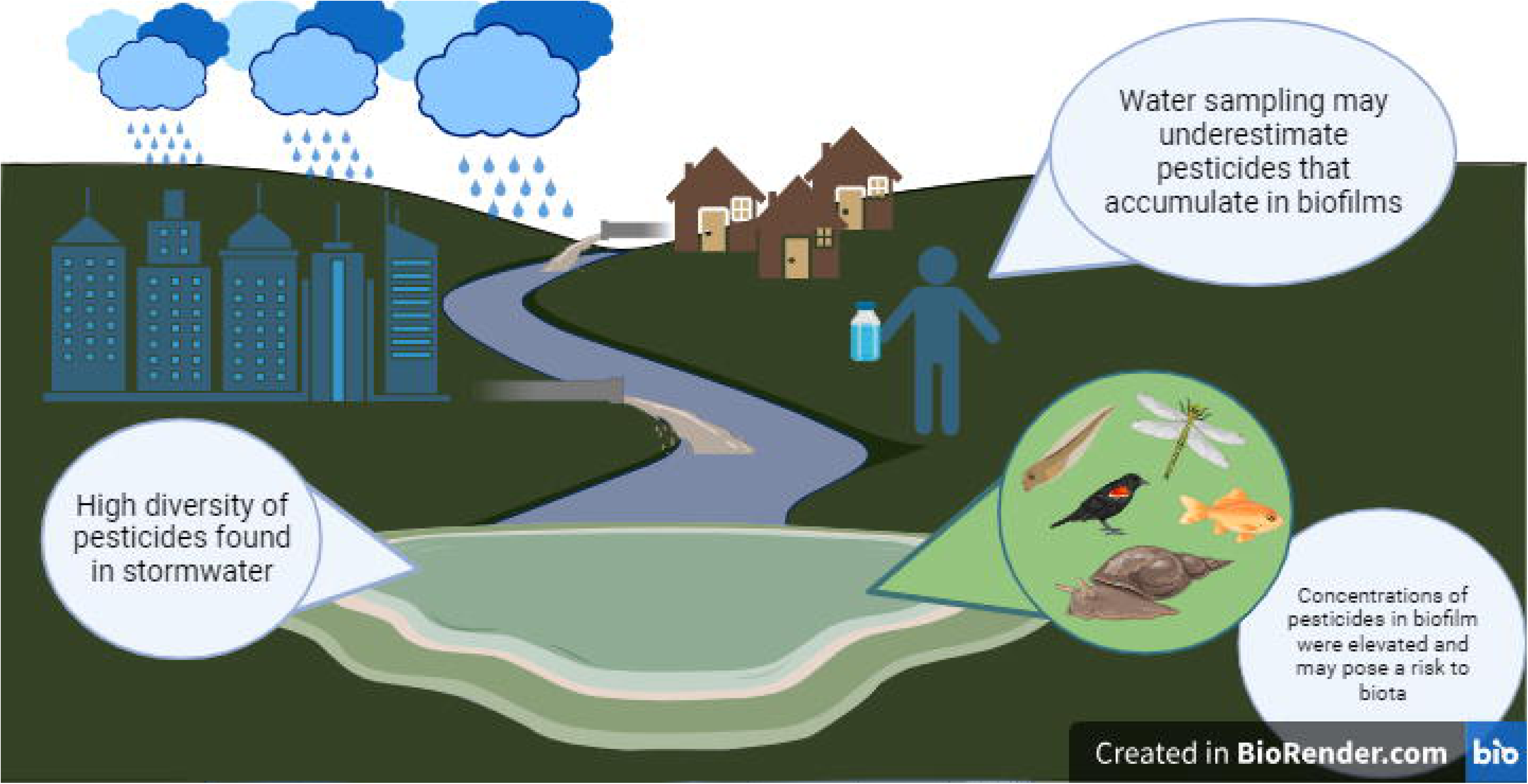

