## Supplemental Information for "Urban-use pesticides in stormwater ponds and their accumulation in biofilms"

*corresponding author

Supplemental Information

Contents:

1. List of pesticide analytes and their MDLs/MQLs in water and biofilm samples
2. Methodology for pesticide analysis
3. Table S1. Water Quality Index Scores
4. Figure S1. Photos of biofilm samplers.
5. Table S2. Pesticide detections in stormwater and in biofilm by pesticide type and use.
6. Pesticide detections and concentrations in water samples
7. Pesticide detections and concentrations in biofilm samples
8. Figure S2. GLMs of pesticide detections in water and biofilm samples.
9. Figure S3. GLMs of pesticide BCFs and physicochemical properties
10. Figure S4. GLM of pesticide BCFs and water quality
11. Figure S5. GLM of pesticide BCFs and ash-free dry weight
12. Other information

**List of pesticide analytes and their MDLs/MQLs in water samples**:
Available on FigShare: <https://figshare.com/s/549156f3317ddc839d69>

**Methodology for pesticide analysis:**

Provided by Brian Atkinson, Agriculture and Food Lab, Guelph, Ontario.

*Subsampling/Preprocessing*

Biofilm samples are homogenized to a powder with dry ice and a small food processor to ensure test portions taken for analysis is representative of the entire sample. For storm water samples are thawed and shaken for one minute by hand before being subsampled for each test. No filtering of the water was used. Meaningful residue data can only be obtained when a representative sample is obtained.

*Sample Analysis*

All sample analyses were performed at the ISO/IEC 17025 accredited Laboratory Service Division, University of Guelph. The multi-residue pesticide determination (AFL Method ID: TOPS-142) was performed by liquid chromatography/electrospray ionization-tandem mass spectrometry (LC/ESI-MS/MS) and gas chromatography-tandem mass spectrometry (GC-MS/MS). Pesticides were extracted from water and biofilm using the method known as the quick, easy, cheap, effective, rugged, and safe (QuEChERS) method with dispersive solid phase extraction (SPE). A representative sample (20mL for water and 3grams for biofilm) was extracted into acetic acid in acetonitrile in the presence of anhydrous sodium acetate and magnesium sulfate. The supernatant was centrifuged and then split, evaporated and diluted with methanol/ammonium acetate (for LC analysis) or hexane (for GC analysis). Sample extracts were analyzed in positive polarity mode using SCIEX5550 ESI-MS/MS with Agilent 1260 HPLC and on Agilent GC 7890 Quadrupole GC-MS/MS. This method detects for over 500 pesticide compounds.

The Phenoxy pesticide screen (AFL Method ID: CHEM-069) was a general solid phase extraction (SPE) 2 step method. 500 mL’s of water is initially acidified with sulfuric acid, passed through the SPE, eluted with methanol, concentrated and analyzed in both positive and negative modes using a SCIEX 5500 LC-MS/MS with an Agilent HPLC. This method screens for 19 phenoxy acidic herbicides as well as for 9 neonicotinoid class pesticides. For the biofilm samples a 3 gram subsample is extracted using a QuEChERS style method of acidified (with formic acid) acetonitrile in the presence of anhydrous magnesium sulfate and citrate buffers. The supernatant centrifuged followed by dilution into a 50:50 MeOH/Water mixture filtered and analyzed in both positive and negative modes using a SCIEX 5500 LC-MS/MS with an Agilent HPLC.

The Polar pesticide screen (AFL Method ID: CHEM-334) is a modified version of the EU’s QuPPE method. A representative subsample (10 mL’s for water and 2 gram for biofilm-w 5 mL’s of Nanopure water to rehydrate) are extracted using acidified (with formic acid) methanol and then placed into a freezer for 30 minutes. The supernatant is then centrifuged and filtered and analyzed in positive and negative modes using a SCIEX 5500 LC-MS/MS with an Agilent HPLC.

*Quality Controls*

All methods use several quality control points that confirm the presence of a compound in the sample: retention time (RT), M+1 target mass, 2 fragment qualifier ions, and the ratio of the two fragment ions. For the identification and quantification of the compounds, each method utilizes deuterium labelled internal standards, matrix matched blanks, spikes, and calibration curves.

**Table S1**. Water Quality Index (WQI) scores for each stormwater pond site calculated using the CCME Water Quality Index Calculator version 1.2.

| Stormwater pond site | CCME WQI |
| --- | --- |
| 130 | 33.5 |
| 109 | 35.6 |
| 116 | 44.7 |
| 26 | 46.2 |
| 153 | 47.2 |
| 191 | 47.5 |
| 106 | 49.4 |
| 51 | 50.7 |
| 174 | 50.8 |
| 10 | 56.8 |
| 49 | 56.9 |
| 46 | 59.2 |
| 58 | 60.1 |
| 188 | 61.2 |
| 56 | 64.6 |
| 14 | 67.1 |
| 85 | 71.5 |
| 96 | 72.2 |
| 113 | 74.0 |
| 156 | 76.1 |
| 78 | 94.6 |

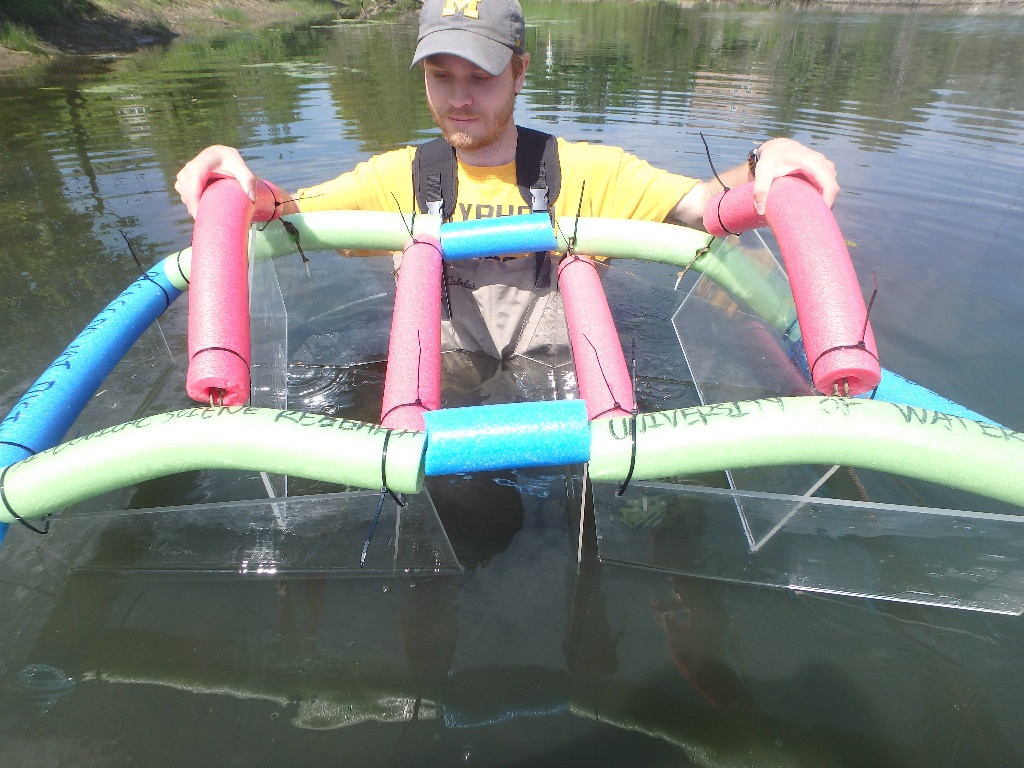

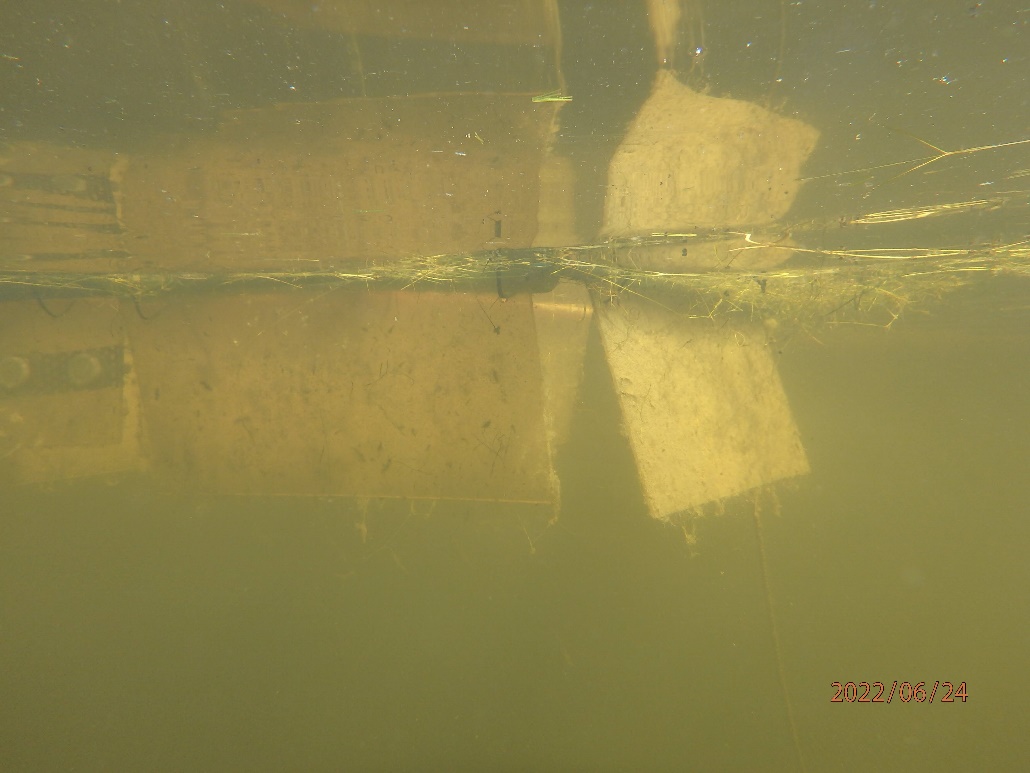

**Figure S1**. Biofilm samplers deployed at stormwater pond sites in Brampton, ON. Acrylic plates were suspended from floats (A) and submerged in the water where they were colonized by biofilms (B).

**Table S2**. Pesticide detections in stormwater and in biofilm by pesticide type and use.

| Category | | Total # of pesticides | Total overall detections | Detections in stormwater | | Detections in biofilm | |
| --- | --- | --- | --- | --- | --- | --- | --- |
|  |  |  |  | # of pesticides | # of hits | # of pesticides | # of hits |
| Type | Herbicides | 14 | 177 | 13 | 139 | 6 | 37 |
|  | Insecticides | 5 | 43 | 5 | 43 | 0 | 0 |
|  | Fungicides | 8 | 62 | 3 | 10 | 7 | 52 |
|  | Other | 5 | 28 | 2 | 13 | 3 | 15 |
| Use | Current-use | 26 | 282 | 21 | 195 | 12 | 87 |
|  | Legacy | 1 | 5 | 1 | 5 | 0 | 0 |
|  | Unknown | 5 | 22 | 1 | 5 | 4 | 17 |

**Pesticide detections and concentrations in water samples:**
Available on Figshare: <https://figshare.com/s/27341527655e5b2b258a>

**Pesticide detections and concentrations in biofilm samples:**
Available on Figshare: <https://figshare.com/s/31e4a23e5819b2517979>

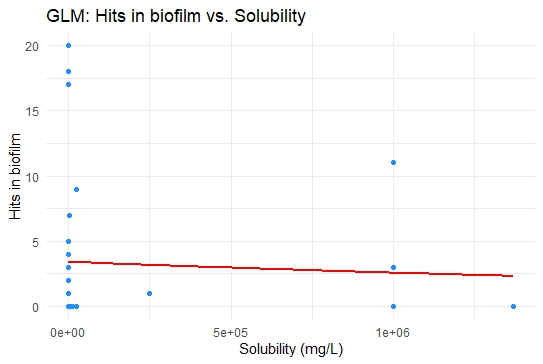

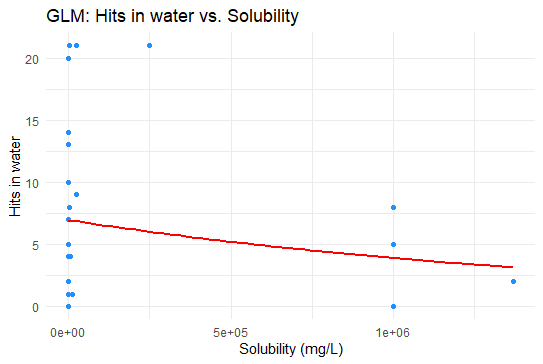

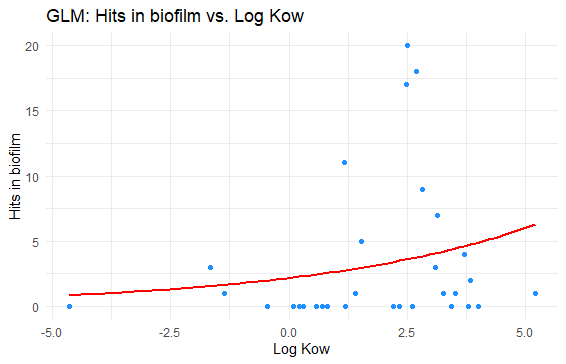

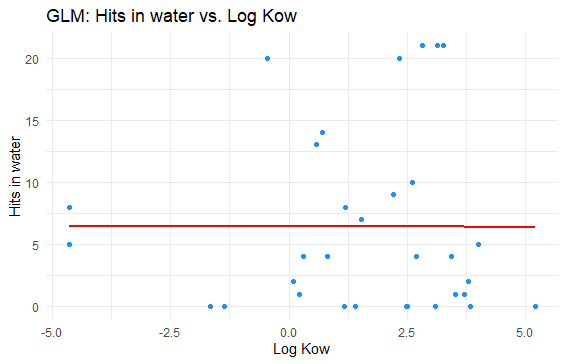

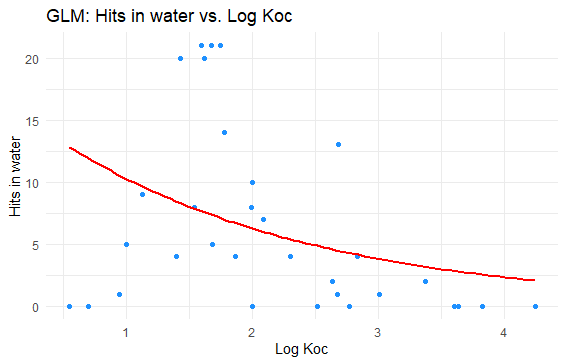

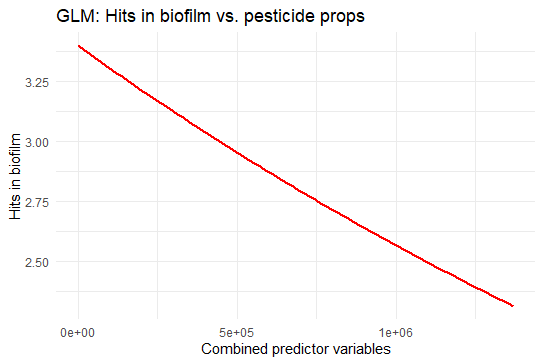

F

E

D

C

B

A

**Figure S2**. Generalized linear models of the solubility (mg L^-1^) of pesticides and their detections (“hits”) in biofilm (A) and water (B); Log K_ow_ of pesticides and their detections in biofilm (C) and water); Log K_oc_ of pesticides and their detections in water (E); and the number of pesticide detections in biofilm using a combined model (F).

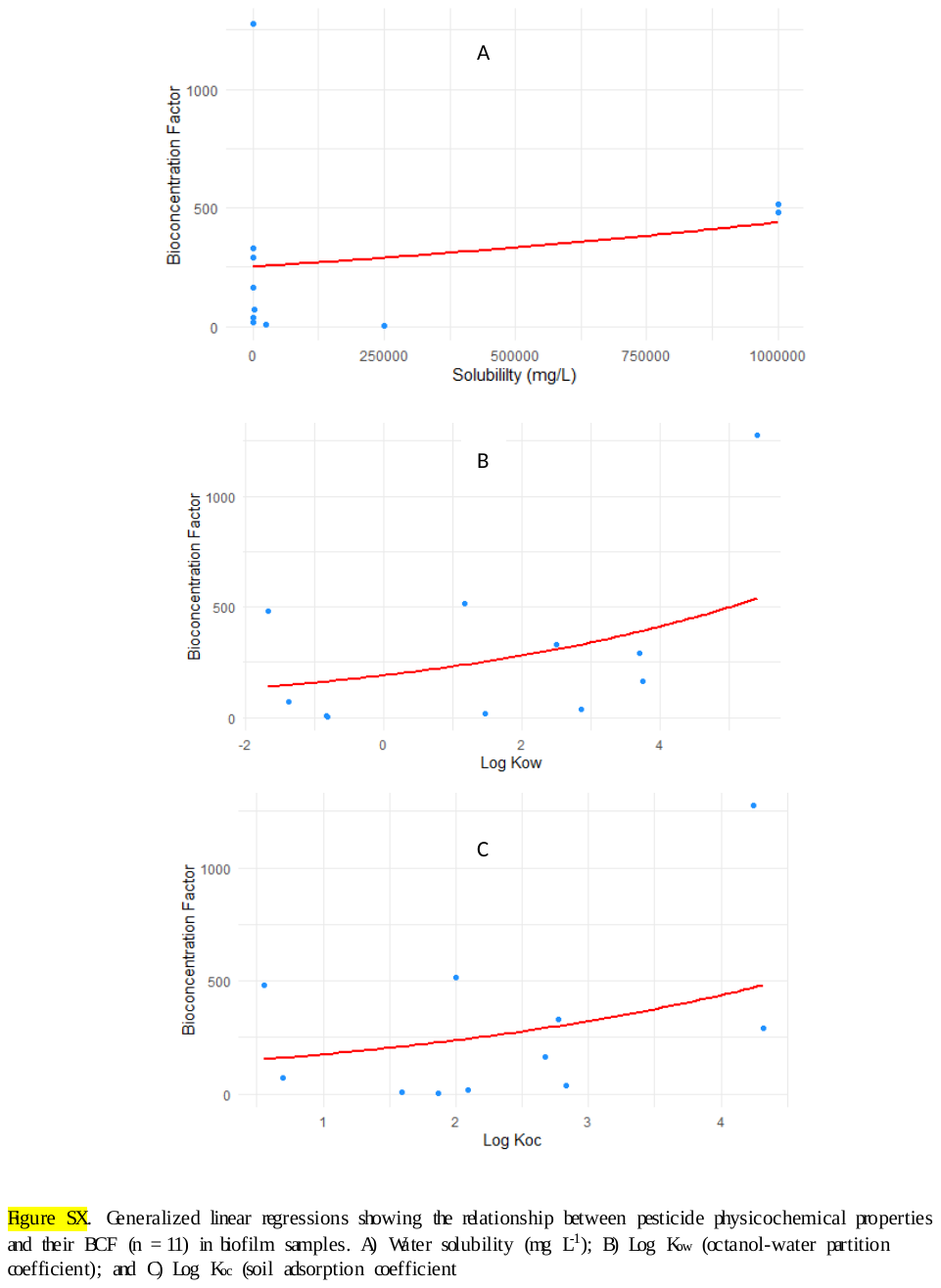

**Figure S3**. Generalized linear regressions showing the relationship between pesticide physicochemical properties and their BCF (n = 11) in biofilm samples. A) Water solubility (mg L-1); B) Log K_ow_ (octanol-water partition coefficient); and C) Log K_oc_ (soil adsorption coefficient)

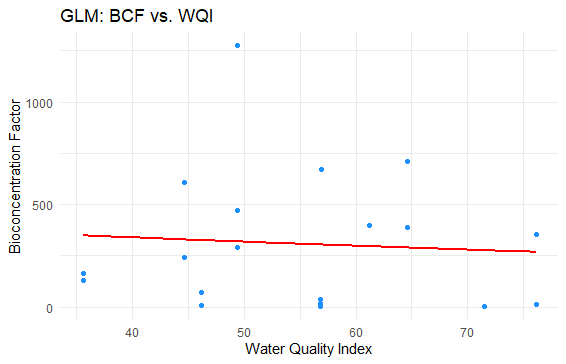

**Figure S4.** Generalized linear model of the CCME Water Quality Index Score and BCF values (n = 17, p = 0.768).

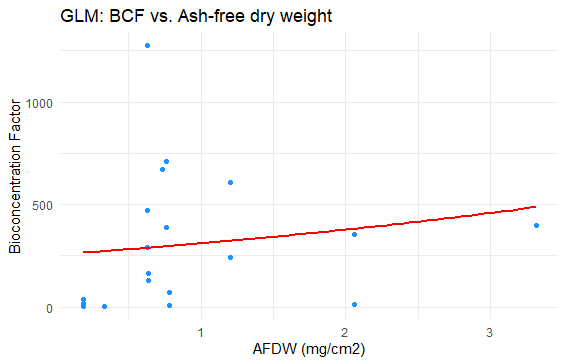

**Figure S5**. Relationship between ash-free dry weight (mg cm^-2^) and calculated pesticide BCF values (n = 17; p = 0.586) using a generalized linear model.

**Other information:**

Site locations: <https://figshare.com/s/8ad9316ca7c42c1928ff>

Site characteristics: <https://figshare.com/s/3f17702070cb8baa26d3>

Water quality assessment results: <https://figshare.com/s/62c747135f94d0672a3d>

E.coli and coliforms (AFL): <https://figshare.com/s/e5cfd674c74bb9aaed70>

Nutrients in water (ALS): <https://figshare.com/s/1bcf2ed9f695a6bdb23d>

CCME WQI Calculations: <https://figshare.com/s/c8aa8abacae79d0ac48c>

Biofilm analysis (chlorophyll-a and AFDW) results: <https://figshare.com/s/76756a987ee8f1590c1c>
